# A generalized growth law for translation- and transcription-targeting antibiotics captures drug interactions

**DOI:** 10.64898/2026.08.21.746212

**Authors:** Natawan Gadjisade, Matteo Mori, Tobias Bollenbach

## Abstract

Bacterial growth laws quantitatively connect intracellular resource allocation to growth rate, enabling accurate predictions of physiology and antibiotic responses. Yet these laws have been rigorously tested for only a handful of perturbations. Here, we show that the growth law linking ribosome levels to growth rate under translation-inhibiting antibiotics is not universal, but rather depends on the antibiotic’s mechanism of action. Quantitative proteomics across finely resolved one- and two-dimensional antibiotic gradients showed that inhibitors of translocation elongation or peptide bond formation elicit the canonical rise in ribosome levels, consistent with the growth law. By contrast, antibiotics disrupting translation initiation or fidelity produced distinct responses without ribosome upregulation. The transcription inhibitor rifampicin even reduced ribosome abundance. Combining antibiotics with divergent ribosome responses revealed a generalized growth law, in which the individual responses to perturbations superimpose. Embedding this law in a mathematical model explains distinct drug interaction patterns observed between rifampicin and different translation inhibitors. A low-dimensional structure pervades the entire proteome, enabling prediction of responses to drug pairs based on single-drug measurements. Together, these findings broaden the scope of bacterial growth laws and provide new principles for predicting responses to antibiotic combinations.

## Introduction

The controlled perturbation of cell physiology using antibiotics has revealed core principles of bacterial systems biology. Bacterial growth laws link exponential growth rate to ribosome concentration, providing a key ingredient for building predictive frameworks for antibiotic responses^1–4^. The ribosome concentration increases linearly with growth rate when nutrient conditions improve, a core principle expressed in the *first growth law*^4–8^. However, under translation inhibition, the ribosome concentration increases linearly as the growth rate is decreased, with a slope that depends on the nutrient environment (*second growth law*)^5^. Remarkably, both relationships are believed to be largely independent of molecular detail, holding across diverse ways of altering nutrient conditions and inhibiting translation. Thus, growth laws have the potential to bridge the gap between physiology and phenotypic plasticity, serving as empirical laws in biology, much like Ohm’s law or the ideal gas law in physics^5,6^.

However, it is unclear how broadly growth laws apply, given that they have mostly been investigated using a limited number of perturbations in *Escherichia coli* laboratory strains^5,9,10^. Translation proceeds through multiple steps, and different translation inhibitors target distinct stages^11,12^. Yet the second growth law has only been rigorously validated for a handful of translation inhibitors—including chloramphenicol, tetracycline, and erythromycin—using the RNA-to-total-protein ratio (RNA/protein) as a proxy for ribosome content^5,9,13^. Recent advances in mass-spectrometry now allow direct quantification of protein abundances^14,15^. Proteomics has revealed how bacteria adjust their proteome allocation beyond the ribosome^16^. Under translation inhibition, for instance, ribosomal protein levels rise, while levels of other proteins, such as enzymes involved in carbon metabolism, fall^14,15^. For chloramphenicol, proteomics data closely align with RNA/protein-based measurements of ribosome levels, and thus with the second growth law^5,9,15^. Therefore, a systematic proteomic comparison across mechanistically distinct translation inhibitors would help assess the generality of the second growth law.

This issue is important for mathematical models based on bacterial growth laws, which can accurately describe bacterial responses to antibiotics^1,9,17^. These models have successfully predicted the dependence of antibiotic susceptibility on nutrient conditions and the shape of the dose-response curve, which relates growth rate to antibiotic concentration^1,17^. The latter enables the inference of binding parameters of translation inhibitors without direct measurement^1^. Notably, bistable regimes, in which bacteria either grow or fail to grow depending on prior history, have been predicted and experimentally validated in different contexts^17–19^. All of these successes are based on one premise: translation inhibitors reduce the number of active ribosomes, prompting the cell to compensate by increasing ribosome synthesis according to the second growth law^1,5,9^.

A closely related mathematical model has been used to predict drug interactions between translation inhibitors^2,3^. Synergistic or antagonistic drug interactions occur when the combined effect of two antibiotics on growth is stronger or weaker than expected based on an additive reference, respectively^20–25^. Antagonistic interactions are common among translation inhibitors because bottlenecks at different stages of translation can partially compensate for one another^2^. Conversely, purely additive interactions are often observed for translation inhibitors with similar modes of action or overlapping ribosomal binding sites^2,24–26^.

In many cases, the individual dose-response curves provide sufficient information to infer uptake and target-binding kinetics, which the model then uses to predict pairwise interactions^2,3,21^. These results show that a quantitative description of global cell physiology can illuminate the mechanistic basis of drug interactions. They also highlight the need to test growth-law predictions in multidrug contexts.

The predictive power of growth laws raises the question of whether analogous relations exist for the perturbation of bacterial functions other than translation. Transcription inhibition is a prime candidate because it affects the synthesis of rRNA, tRNA, and mRNA, all of which are tightly linked to translation^27^. This involvement could be captured by generalizing the existing theoretical framework for translation inhibitors^1,3^. Recent mathematical models have incorporated the role of transcription by considering RNA polymerase (RNAP) and mRNA levels^28–30^. These models predict that growth can sometimes be limited by mRNA availability rather than by ribosomes^30,31^. Another prediction is that ribosome levels can increase, decrease, or remain constant under transcription inhibition depending on the nutrient environment; an increase is predicted in poor nutrient conditions and a decrease in rich conditions^30^. However, experimental validation is sparse: direct measurements of ribosome levels and global proteome allocation under transcription inhibition—alone or combined with translation inhibitors—are missing. For rifampicin, RNA/protein measurements hint at a distinct resource allocation pattern^30,32–34^, but quantitative proteomics under transcription inhibition has yet to be performed.

Beyond ribosome levels, global responses to drug combinations may follow general principles^35–37^, pointing to a more general, low-dimensional structure in how biological systems respond to perturbations^38^. High-resolution transcriptional measurements across two-dimensional drug concentration gradients revealed that most gene expression levels interpolate the responses to the individual drugs along lines of constant growth rate in both *E. coli*^36^ and yeast^37^. Depending on the drug interaction, the transition can be either smooth and continuous or relatively abrupt^35–37^. This growth-rate-anchored interpolation is a candidate empirical law for predicting the effects of combinations from single-drug data, but it has not yet been tested at the protein level.

Here, we report how translation and transcription inhibitors differentially impact *E. coli* proteome allocation, and how these differences shape drug interactions. We found that certain translation inhibitors violate the second growth law, as ribosome concentration remained constant despite a decrease in growth rate. The transcription inhibitor rifampicin even induced a linear decline in ribosome levels. Combining translation inhibitors with rifampicin revealed a generalized growth law that partly explains the differences in drug interactions between rifampicin and different translation inhibitors. High-resolution proteomics across two-dimensional drug gradients further revealed a low-dimensional structure of the response space, enabling partial prediction of proteome reallocation and drug interaction under multidrug stress.

## Results

### Ribosome responses to antibiotics depend on drug mode of action

To quantify how the bacterial proteome reorganizes in response to antibiotics, we performed quantitative data-independent acquisition (DIA) mass-spectrometry proteomics on *E. coli* cultures in steady-state exponential growth exposed to various translation and transcription inhibitors. We inferred protein abundances from peptide-level measurements using the xTop algorithm and converted them to absolute proteome mass fractions with the help of published ribosome-profiling reference data^39^ (Methods). We collected two independent datasets under different cultivation conditions: one in a mini-bioreactor (15 mL) under tightly controlled growth at fixed optical density, and another in flasks where cultures were precultured and diluted into antibiotic-containing medium (20 mL). Both ensured balanced exponential growth at the time of sampling, as verified by optical density tracking (Methods). We generated a comprehensive dataset of several conditions in duplicate using both techniques. We profiled proteomes across graded growth inhibition induced by carbon limitation, as well as one- and two-dimensional gradients of antibiotics targeting different stages of translation and transcription (Fig. 1A). Across both datasets, 1,700 to 2,400 proteins were detected per condition, including at least 43 ribosomal proteins and at least 7 RNAP-associated proteins. Reproducibility was high across both replicates and cultivation procedures (Methods). The resulting dataset provides a high-resolution map of proteome allocation under various stressors.

**Fig. 1:**
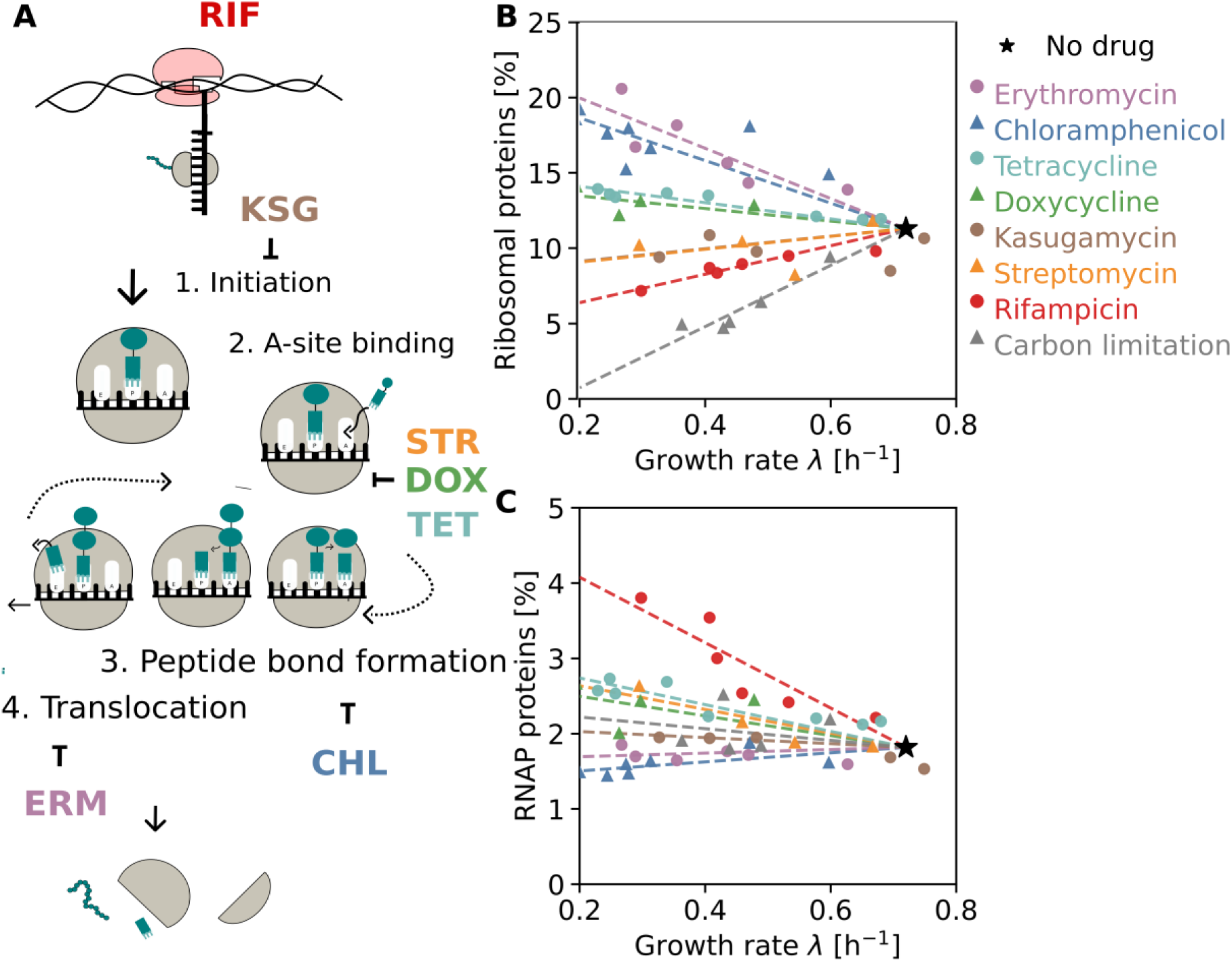
Quantitative proteomics reveals drug-specific ribosome-growth relationships for translation- and transcription-targeting antibiotics. **A)** Schematic overview of the primary molecular targets of the antibiotics used in this study, including diverse translation inhibitors (streptomycin, STR; doxycycline, DOX; tetracycline, TET; erythromycin, ERM; chloramphenicol, CHL, kasugamycin, KSG) and the transcription inhibitor rifampicin (RIF). **B)** Ribosomal protein mass fraction as a function of growth rate under different antibiotics (colors) or carbon limitation (gray; Methods). **C)** As B, but for the RNA polymerase-associated protein mass fraction (Methods). Data in B-C were obtained from bioreactor-grown cultures (Methods).

Our proteomics experiments reproduced the canonical growth-law relationships for ribosome allocation. To determine how these changes in ribosome levels were accompanied by broader proteome reorganization, we assigned proteins to functional sectors^40^ and tracked the proteome mass fractions of different sectors across conditions and growth rates (Table S1). As a physiological benchmark, we first examined carbon limitation induced by increasing concentrations of α-methylglucoside, a non-metabolizable glucose analog (Methods). Under this perturbation, growth rate and ribosome concentration decreased in parallel, following an approximately linear relationship, consistent with the first growth law^9^ (Fig. 1B). In contrast, the translation inhibitors chloramphenicol and erythromycin, which block peptide bond formation and translocation, respectively (Fig. 1A), caused an approximately linear increase in ribosome concentration as growth rate decreased (Fig. 1B), in line with the second growth law^9^. This ribosome upregulation was accompanied by proportional downregulation of other sectors, particularly central carbon metabolism (Fig. S1; Table S1)^14,15^, consistent with a feedback loop in which stalled ribosomes stimulate the production of more ribosomes to maintain translational capacity^1,9,29,41^. The agreement under previously reported perturbations validates the quantitative accuracy of our proteomic measurements^14,15^.

Motivated by numerous mathematical models of bacterial translation inhibition that invoke the compensatory ribosome upregulation captured by the second growth law^1–3,5,9,17^, we asked whether this response is conserved across antibiotics that target different stages of protein synthesis. Tetracycline and doxycycline, which block tRNA entry into the ribosomal A-site^42^, induced an increase in ribosome concentration. However, this increase was modest compared with chloramphenicol and erythromycin (Figs. 1B and S2A). Notably, streptomycin, an aminoglycoside that disrupts translation fidelity^11,12,43^, and kasugamycin, which blocks translation initiation^44^, failed to elicit ribosome upregulation (Figs. 1B and S2A) and produced only modest sector-level reorganization relative to untreated bacteria (Table S1). Recent RNA/protein measurements revealed a similar lack of ribosome upregulation when translation initiation was inhibited using CRISPRi to knock down translation initiation factor IF-1 (*infA*)^45^. These differences suggest that ribosome compensation is most effective when translation elongation is slowed or stalled, allowing bacteria to offset reduced translational throughput by increasing ribosome abundance. In contrast, the effects of antibiotics that block initiation or corrupt decoding fidelity appear to be compensated for less effectively^11,12,42,43^. Together, these data show that there is no universal proteome response to translation inhibition; instead, the response reflects the specific molecular mechanism of different antibiotics.

Next, we explored how the proteome adapts to transcriptional inhibition, a fundamentally different perturbation that indirectly impairs translation by reducing the supply of rRNAs, mRNAs, and tRNAs. Unlike translation inhibitors, rifampicin, an antibiotic that blocks transcription initiation by inhibiting RNAP^32^, caused a decline in ribosome concentration as growth rate decreased, following an approximately linear trend, albeit less steep than under carbon limitation (Fig. 1B). In parallel, the mass fraction of RNAP increased substantially in response to rifampicin, again following an approximately linear trend (Fig. 1C; Methods). This reciprocal response – ribosome levels down, RNAP levels up – mirrors the ribosome upregulation observed under translation inhibition (Fig. 1B), but in reverse: here, the bacteria appear to boost the abundance of the inhibited enzyme to counteract reduced transcriptional flux. In line with this view, RNAP levels were essentially unchanged under most translation inhibitors, showing only minor increases, e.g., under tetracycline, and a slight decrease under chloramphenicol (Fig. 1C). We also observed that central carbon pathways were expressed at higher levels in rifampicin-treated cells than in translation-inhibited cells, and at similar levels to those in carbon-starved cells (Fig. S1). Notably, the expression level of the premature transcription termination factor Rho, which plays a key role in coordinating transcription-translation coupling^46,47^, behaved similarly to ribosomal protein concentration across all perturbations investigated here (Fig. S2C). This suggests that the need for premature transcription termination changes in tandem with total translation capacity. Together, these results show that antibiotics elicit distinct proteome allocation responses depending on whether translation or transcription is perturbed. They also highlight that core cellular machinery, such as the ribosome, can be regulated in opposite ways by closely related perturbations.

### Combining transcription and translation inhibitors reveals a generalized growth law

The contrasting responses to translational and transcriptional inhibition raise the question of how bacteria respond when both processes are inhibited simultaneously: Do the combined perturbations generate a qualitatively different proteome allocation state, or do bacteria transition continuously between the single-drug responses corresponding to each process? To resolve this, we systematically mapped proteome reorganization under combined antibiotic perturbations, starting with ribosomal proteins and transcriptional machinery, and then extending the analysis to the entire proteome.

We first examined the combination of rifampicin and chloramphenicol, which elicit strongly opposing ribosomal allocation responses when applied alone (Fig. 1B). Ribosome allocation followed a superposition of the responses to the individual drugs. Progressively increasing chloramphenicol concentrations at a fixed rifampicin concentration resulted in an upregulation of ribosome levels, following an approximately linear trend (Fig. 2A,B). The slope of this increase increased with decreasing growth rate in the rifampicin-only condition (Fig. 2B). Notably, this behavior resembles that observed across nutrient conditions in the second growth law^5,7–9^, despite quantitative differences. Conversely, increasing the rifampicin concentration at fixed chloramphenicol concentration pulled the system to the lower-ribosome rifampicin state (Fig. S3A,B). Independent measurements under balanced exponential growth conditions in a controlled bioreactor reproduced the observed growth-ribosome trajectories (Fig. S3C,D; Methods). Therefore, the combined rifampicin– chloramphenicol treatment robustly produced a superposition of the two opposing ribosome responses to the individual drugs.

**Fig. 2:**
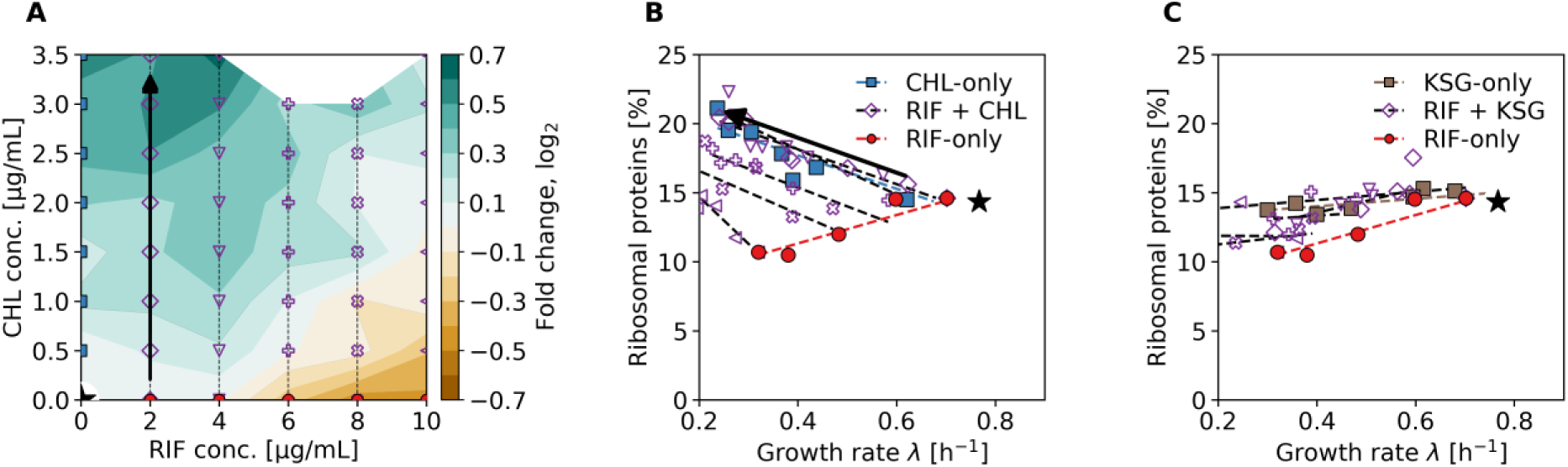
Ribosome response to antibiotic combinations is a superposition of the responses to the individual drugs, suggesting a generalized growth law. **A)** Heatmap of ribosomal protein abundance relative to the no-drug condition across a two-dimensional concentration gradient of rifampicin (RIF) and chloramphenicol (CHL). Black arrow indicates a trajectory of increasing chloramphenicol concentration at one fixed rifampicin level. **B)** Ribosomal protein mass fraction as a function of growth rate for rifampicin-only, chloramphenicol-only, and combined rifampicin-chloramphenicol conditions at the concentrations indicated by the same symbols in A. The combined response is well approximated by a superposition of the two single-drug responses. **C)** As B, but for the combination of rifampicin and kasugamycin (KSG). Data in A-C were obtained from cultures grown in flasks (Methods).

To test whether this superposition principle holds more generally, we examined rifampicin combined with kasugamycin—a drug that, on its own, leaves ribosome levels essentially unchanged (Fig. 1B). The corresponding ribosome concentrations remained tightly constrained across conditions (Fig. 2C). Increasing the kasugamycin concentration at a fixed rifampicin concentration caused only modest deviations from the rifampicin trajectory (Fig. 2C), reflecting the limited response induced by kasugamycin alone. As with the rifampicin– chloramphenicol pair, the combined response was consistent with a superposition of the single-drug responses, albeit within a narrow dynamic range that precluded detailed quantitative analysis.

These results suggest that, under combined drug stress, ribosome allocation follows a generalized growth law in which the two single-drug states are superimposed (Fig. 2B,C). They further imply that mathematical models of bacterial physiology based on growth laws^1–3^, which connect cellular resource allocation to fitness, can be extended rigorously to antibiotic combinations, when relaxing the ad hoc assumption that the second growth law holds universally^1–3,21^.

### Antibiotic-specific ribosome responses rescale drug susceptibility and reshape drug interactions

We asked whether the drug-specific ribosome responses (Fig. 1B) could account for the differences in drug interactions observed when different translation inhibitors are combined with a transcription inhibitor. To characterize these interactions, we mapped lines of constant growth rate (isoboles) in two-dimensional drug space, using Loewe additivity as the null model^20^. Straight isoboles indicate additivity; convex isoboles indicate synergy (i.e., stronger-than-additive growth inhibition); and concave isoboles indicate antagonism (i.e., weaker-than-additive growth inhibition) (Fig. 3A). To quantify global drug interaction patterns using a compact metric, we computed the interaction score, LI_log_, which is defined as the logarithm of the ratio of the integrated measured growth response surface to the corresponding additive reference surface^2^. Positive LI_log_ values indicate antagonism, LI_log_ ≈ 0 indicates additivity, and negative values indicate synergy (Methods). The strongest interactions between ribosome inhibitors, which include highly suppressive interactions caused by ribosome traffic jams on mRNAs, have |LI_log_| > 0.2, but interactions with |LI_log_| > 0.1 are robustly detectable^2^.

**Fig. 3:**
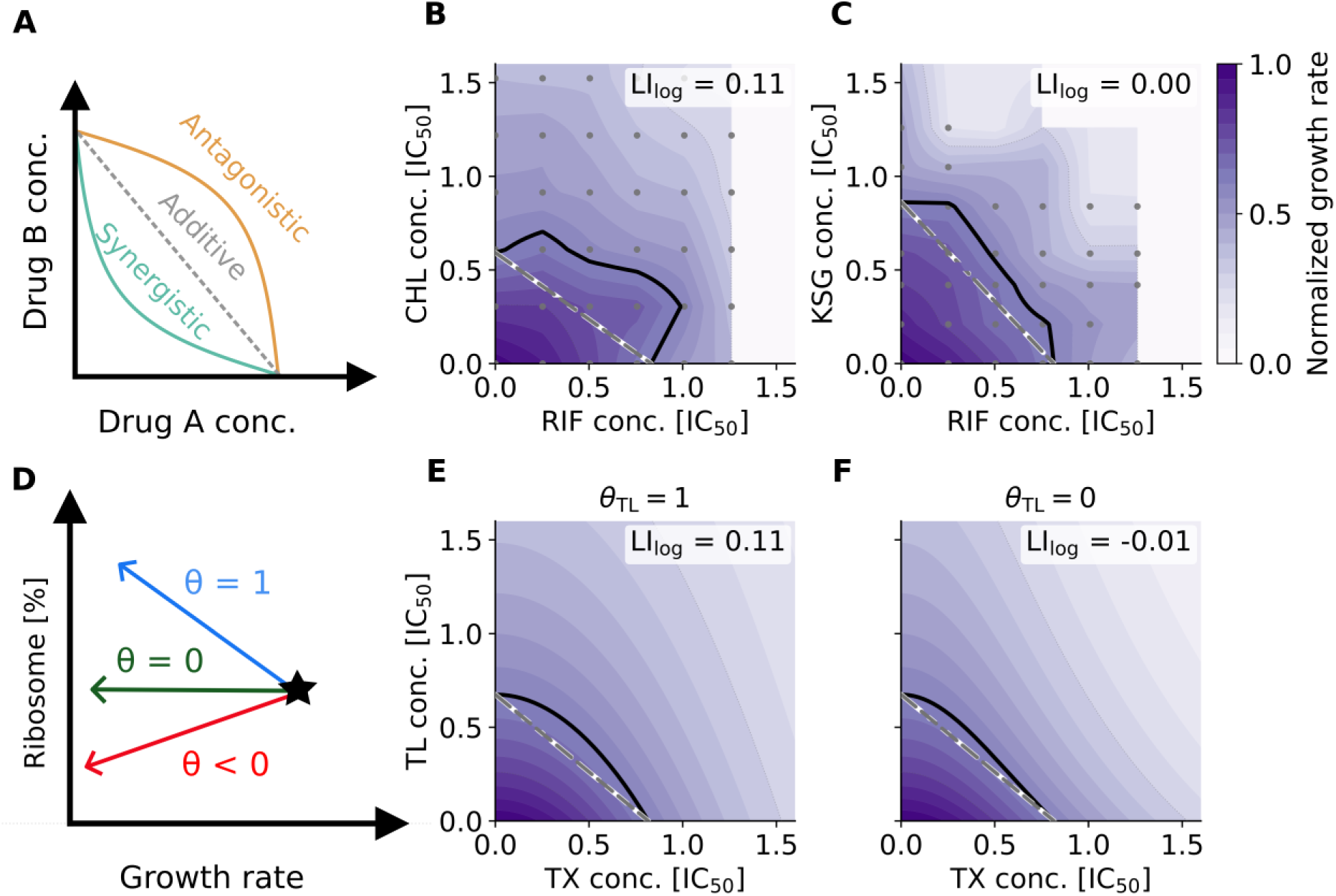
A mathematical model based on the generalized growth law captures much of the observed variation in drug interactions. **A)** Schematic of Loewe additivity in two-drug concentration space^20^. Lines of constant growth rate (isoboles) are shown. Straight isoboles (gray) indicate additive interactions. Concave (orange) and convex isoboles (green) indicate antagonistic and synergistic interactions, respectively. **B)** Growth-rate landscape of rifampicin (RIF) and chloramphenicol (CHL) combinations in two-dimensional concentration space. The isobole at 40% growth inhibition (black line) is concave, indicating strong antagonism. The interaction score, LI_log_=0.11, quantifies the strength of the drug interaction based on the entire landscape (Methods); LI_log_>0 for antagonism and LI_log_<0 for synergy. Gray points indicate the experimentally measured conditions; intermediate regions were interpolated (Methods). **C)** As B, but for the combination of rifampicin and kasugamycin (KSG). The isobole remains close to the Loewe-additive expectation, indicating near-additive behavior, consistent with interaction score LI_log_≈0. **D)** Schematic representation of growth laws with different ribosome-response parameters (slopes) *θ* for different antibiotics targeting translation and transcription (Supplementary Text). Some translation inhibitors elicit strongly increasing ribosome allocation (*θ*=1), others weak ribosome responses (*θ*=0); transcription inhibition is captured by *θ*<0. **E)** Growth-rate landscape for combination of a transcription inhibitor (TX) and a translation inhibitor (TL) that elicits strong ribosome upregulation (*θ*=1), calculated from a mathematical model of bacterial growth (Methods). The model produces a clearly antagonistic interaction (LI_log_=0.11), as observed experimentally (B). **F)** As E, but for a translation inhibitor with a weak ribosome responses (*θ*=0). The model produces a nearly additive interaction (LI_log_=-0.01), as observed experimentally (C). Data in B-C were obtained from cultures grown in flasks (Methods).

Although both drugs target translation, chloramphenicol and kasugamycin behaved differently when paired with the transcription inhibitor rifampicin. The rifampicin– chloramphenicol growth response surface revealed pronounced antagonism between the two antibiotics (LI_log_ = 0.11; Fig. 3B), a result that was independently reproduced in a mini-bioreactor (LI_log_ = 0.12; Fig. S4B). By contrast, the rifampicin–kasugamycin growth response surface remained close to additive (LI_log_ ≈ 0; Fig. 3C). This implies that the combination with compatible ribosome responses (Fig. 2C) remained additive, while the combination with divergent single-drug responses (Fig. 2B) produced strong antagonism. This observation suggests that ribosome allocation under stress can influence the effects of drug combinations.

To address whether the transition from near-additivity to antagonism between rifampicin and translation inhibitors stems from their distinct ribosome responses, we extended a mathematical model of antibiotic effects on bacterial growth. This extension relaxes the fixed ribosome-growth relationship (second growth law) and introduces a drug-specific ribosome-response parameter, *θ*, which scales the magnitude of ribosomal up- or down-regulation (Supplementary Text). We analyzed the model in steady state, corresponding to balanced exponential growth at fixed drug concentrations. In this framework, *θ* = 1 reproduces the strong compensatory ribosome upregulation of the second growth law observed with chloramphenicol, *θ* ≈ 0 captures the minimal reallocation as observed with kasugamycin, and *θ* < 0 describes the ribosome downregulation induced by rifampicin (Fig. 1B). The resulting generalized ribosome-growth relation (Fig. 3D), which extends previously used growth laws^1–3,5^, is given by

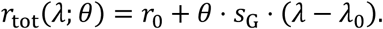

Where *r*_tot_ is the total ribosome concentration, *λ* is the growth rate, *r*_0_ and *λ*_0_ denote the ribosome concentration and growth rate in the drug-free reference condition, respectively, and *s*_G_ is the slope of the original second growth law^5^. This generalization captures the spectrum of experimentally observed ribosome responses while maintaining the predictive power of earlier growth-law models.

In this model, modulating the ribosome response preserves the intrinsic shape of the dose-response curve for translation inhibitors, but rescales the effective drug concentration axis, resulting in a horizontal shift of the curve (Fig. S4A). Thus, the ribosome-response parameter *θ* acts as an independent lever on drug susceptibility. In particular, a lack of compensatory ribosome upregulation (*θ* ≈ 0) markedly increases susceptibility to translation inhibitors (Fig. S4A). By contrast, altering the shape of the dose-response curve requires changes to kinetic parameters, such as drug binding, unbinding, or transport^1^. This finding highlights the ribosome-response parameter *θ* as a key regulatory node that modulates antibiotic susceptibility without altering their intrinsic potency.

To capture the joint action of a transcription inhibitor and a translation inhibitor, we next extended the model to two drugs. We treated the transcription inhibitor (drug X, e.g., rifampicin) as a modifier of the physiological background, while the translation inhibitor (drug Y) acts within this altered background. Based on our experimental observations (Figs. 1B and 2B-C), we described the transcription inhibitor as reducing the effective growth rate in a manner analogous to nutrient limitation:

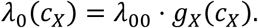

Here, *g_X_*(*c_X_*) is the normalized dose-response curve of drug X, *λ*_00_ denotes the drug-free growth rate, and *λ*_0_(*c_X_*) is the background growth rate established by drug X. Thus, the transcription inhibitor shifts the physiological context in which the translation inhibitor operates.

We assumed that the ribosome response to the translation inhibitor retains the same functional form as in the single-drug case, but is applied to the physiological state set by the transcription inhibitor (Supplementary text). Consequently, both the dynamic range of accessible ribosome concentrations and the susceptibility to the translation inhibitor become functions of the transcription inhibitor concentration, matching the trends seen experimentally (Fig. 2B,C). Solving the model showed that, at each fixed transcription inhibitor concentration, the dose-response curve of the translation inhibitor is simply rescaled (Fig. S4A), and the system remains monostable. Therefore, the steady state reached is independent of the order in which the drugs are added. The full set of these rescaled dose-response curves constitutes the growth response surface (Supplementary text). This two-drug model thus reveals how the interaction between transcription and translation inhibitors is governed by the ribosome-response parameter *θ*, provided that the antibiotics do not disrupt other cellular processes (e.g., drug uptake) that are not included in the model.

In contrast to the single-drug case, where different ribosome responses merely rescale the effective dose, in the two-drug case these responses can reshape the interaction landscape. The resulting growth-response surfaces revealed that drug pairs with opposing ribosome responses, such as rifampicin and chloramphenicol, produce markedly concave isoboles, indicating antagonism (Fig. 3E). Conversely, combinations of drugs with weaker, more similar ribosome responses, such as rifampicin and kasugamycin, produce nearly straight isoboles, indicating additivity (Fig. 3F). Thus, the model predicts stronger antagonism for rifampicin–chloramphenicol than for rifampicin–kasugamycin (Fig. 3E,F), a trend that aligns qualitatively with experimental data (Fig. 3B,C). Moreover, the interaction scores closely match, while the detailed shapes of the response surfaces differ, partly due to measurement noise (Fig. 3B,C,E,F). Together, these results provide evidence that drug-specific ribosome responses are key determinants of drug interaction outcomes. More broadly, they reveal how antibiotic-induced proteome reorganization can shape response surfaces, providing a mechanistic link between cellular physiology and drug interactions.

### An interpolation principle captures the proteome-wide response to antibiotic combinations

After finding that the ribosome concentration responds to antibiotic combinations according to a generalized growth law (Fig. 2), we wondered whether a similar principle might govern individual proteins across the entire proteome. Our high-resolution quantitative proteomics dataset spanning fine-grained two-dimensional antibiotic concentration gradients (Figs. 2 and S3; Table S2) enables us to address this question systematically. To dissect how *E. coli* rearranges its proteome globally, we performed principal component analysis (PCA) on the measured protein expression levels (Methods). In essence, this approach extracts the dominant response modes, or principal components (PCs), that capture most of the variance in the data. Each protein’s expression across the entire two-drug space can then be approximated as a linear combination of a small number of PCs (Methods). Previous studies that applied this analysis to transcriptional regulation data under drug combinations in *E. coli* and yeast showed that the first three PCs explain more than 95% of the variance^36,37,48^, revealing a low-dimensional structure that enables predictions of responses to combinatorial perturbations. Here, we extend this framework to the protein level—arguably the most functionally relevant layer of cellular regulation^5,9,14,15^.

The leading PCs dominate the variance in protein levels in response to antibiotic combinations. Across all datasets, the top three PCs collectively account for roughly 80% of the variance (Fig. 4A). However, the explanatory power of additional PCs diminishes rapidly: explaining 95% of the variance typically requires about ten PCs, with components beyond the third each explaining less than 5% (Fig. 4A). This indicates that protein-level responses to drug combinations occupy a higher-dimensional space than transcriptional responses. Still, the first two PCs explain most of the variance, raising the question of whether they reflect biologically interpretable response modes.

**Fig. 4:**
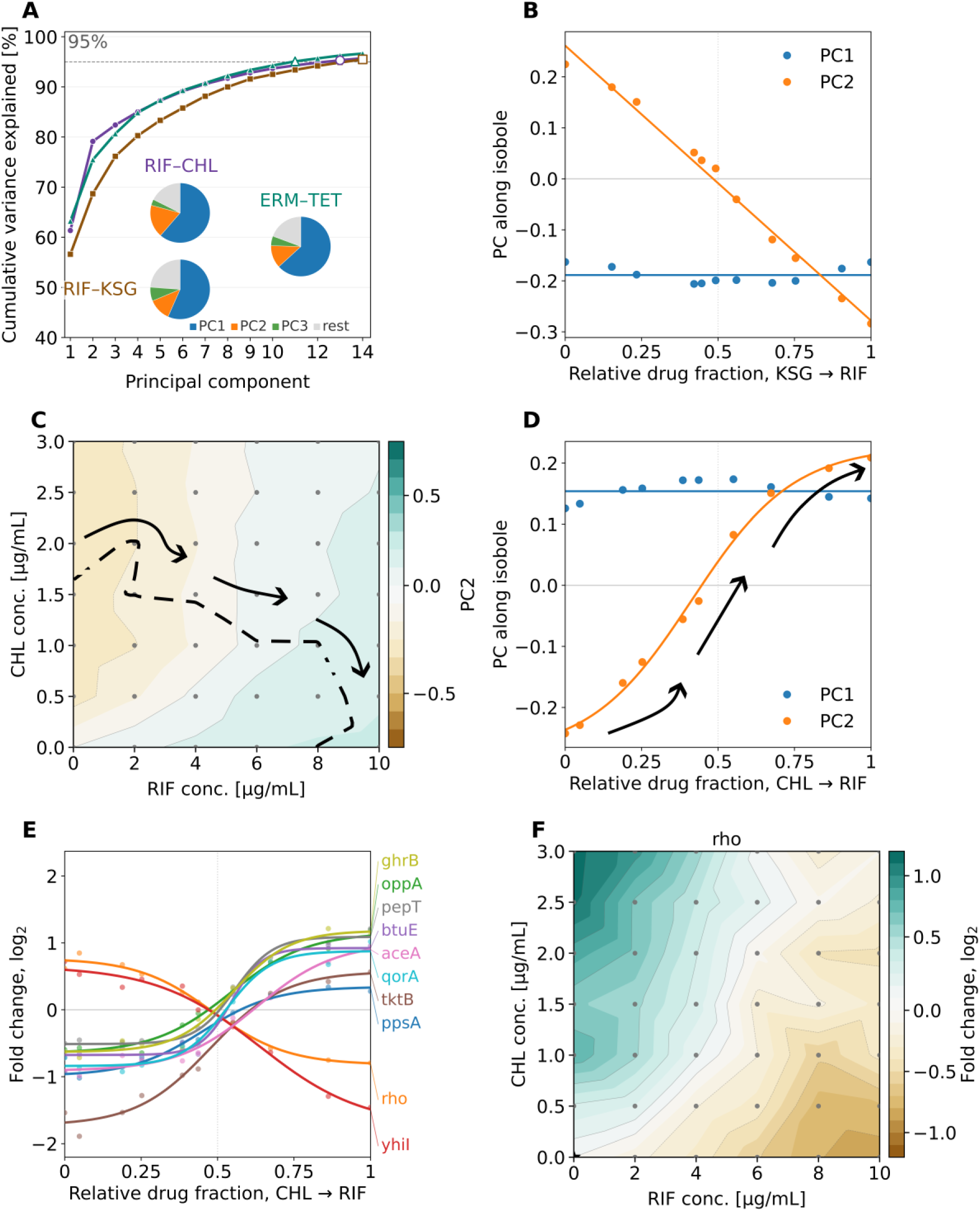
Proteome response to drug combinations is captured by an interpolation of the individual drug responses. **A)** Cumulative fraction of variance explained by principal component analysis (PCA) of proteome responses under the antibiotic combinations rifampicin-chloramphenicol (RIF–CHL, purple), rifampicin-kasugamycin (RIF–KSG, brown), and erythromycin-tetracycline (ERM– TET, green). The pie charts show the variance explained by the first three PCs for each dataset. **B)** Principal components along the isobole at 50% growth inhibition as a function of the relative drug fraction for the near-additive rifampicin-kasugamycin combination (Methods). The first PC remains approximately constant along the isobole, indicating a general growth-related response. The second PC linearly connects the two single-drug responses, indicating gradual interpolation of the conflicting proteome responses. **C)** Second PC for the rifampicin-chloramphenicol combination. Dashed line shows isobole at 50% growth inhibition. **D)** As B, but for the antagonistic rifampicin-chloramphenicol combination. The second PC exhibits a sigmoidal shape, indicating a more switch-like transition between the single-drug states. **E)** Fold-change in expression level of representative proteins along the isobole at 50% growth inhibition for the antagonistic rifampicin-chloramphenicol combination. See Table S2 for the full dataset. **F)** Fold-change in expression level of the premature transcription termination factor Rho in response to the rifampicin-chloramphenicol combination. Additional examples of individual protein responses with clear regulatory conflicts are shown in Fig. S8A-C. Data in A-F were obtained from cultures grown in flasks (Methods).

The first two PCs capture the bulk of the proteome response: a global allocation shift driven by changes in growth rate and a finer adjustment that resolves drug-induced regulatory conflicts. Since growth rate exerts a dominant effect on cellular physiology at all levels^5,7,9,14,15,28,41,46^, we dissected protein levels along isoboles to separate this strong growth effect from other response modes^36,37^. The first PC, which explains 55-65% of the variance for different antibiotic pairs (Fig. 4A), is essentially flat along isoboles (Figs. 4B,D and S5F). In addition, the scores of the first PC across samples were strongly correlated with the corresponding growth rates; the correlation was much weaker when the first PC scores were compared to the relative drug fraction (Fig. S6A-D; Methods). Thus, this component captures the global response to growth-rate modulation, regardless of which drug causes it. The high fraction of variance explained aligns with the expected dominant effect of growth-rate changes on proteome rearrangements. Most of the contribution to the first PC loadings derived from amino acid and nucleotide biosynthetic pathways, which are typically regulated in response to the demand flux via feedback inhibition (Fig. S6E-H; Methods).

The scores of the second PC, which contributes 12-22% of the variance, were not correlated with growth, but rather with the relative drug fraction (Fig. S7A-D; Methods). Thus, the second PC captures how regulatory conflicts are reconciled under drug combinations when protein levels diverge between single-antibiotic treatments. An analysis of how different protein functional groups contribute to the loadings of the second PC highlighted the main changes in the proteome (Fig. S7E-H; Methods). For the rifampicin-chloramphenicol pair, ribosomal proteins had a negative projection, and central carbon pathways had a positive projection (Fig. S7E,F), in agreement with the single drug proteomics results (Fig. 1B, Fig. S1). For this drug pair, we also observed a negative contribution for nucleotide biosynthetic pathways (Fig. S7E,F), possibly due to lower nucleotide consumption in transcription-inhibited cells. For the rifampicin-kasugamycin pair, the results were similar, except for a less prominent contribution from ribosomal proteins (Fig. S7G), due to their more similar growth-rate dependence (Fig. 1B). Finally, the results for the erythromycin-tetracycline pair did not highlight any specific protein groups other than ribosomal proteins (Fig. S7H), indicating that these two drugs mostly elicit responses that differ on a finer scale. Along isoboles traversing the two-drug space, conflicting protein levels are smoothly interpolated (Figs. 4B-D and S5F), as previously observed at the transcriptional level^36,37,48^.

The quantitative pattern of regulatory conflict interpolation mirrors the drug interactions. We used the relative drug fraction, a coordinate derived from each drug’s concentration in the combination relative to its half-maximal inhibitory concentration (IC₅₀), to specify positions along isoboles in two-drug space (Methods). Along the 50% growth-inhibition isobole, the relative drug fraction ranges from 0 (only drug X) to 0.5 (equal contributions of X and Y, relative to their IC₅₀) to 1 (only drug Y). For the nearly additive rifampicin–kasugamycin pair (Fig. 3C), the second PC exhibited an almost perfectly linear interpolation of regulatory conflicts along the isobole (Fig. 4B). A similar linear interpolation emerged for the additive combination of the two translation inhibitors erythromycin and tetracycline (Fig. S5F). By contrast, the antagonistic rifampicin–chloramphenicol pair (Fig. 3B) produced a mildly sigmoidal second PC centered at a relative drug fraction of 0.5 (Fig. 4D), indicating a more switch-like transition between the conflicting single-drug responses.

This interpolation principle was also evident at the level of individual proteins. We classified each protein’s abundance trajectory along IC_50_-normalized isoboles as linear, sigmoidal, or ambiguous based on whether they were better fit by a sigmoidal function or a straight line, while penalizing additional fit parameters (Methods). For the antagonistic rifampicin– chloramphenicol combination, the vast majority (>80%) of classified proteins exhibited switch-like sigmoidal transitions centered at a relative drug fraction of 0.5, rather than gradual linear behavior (Fig. 4E). This points to a prioritization strategy for antagonistic drug pairs, in which bacteria switch from one response program to another, depending on which antibiotic dominates. A notable example is the transcription-termination factor Rho, which transitions relatively abruptly from an elevated expression level under chloramphenicol to a lower level under rifampicin as the relative drug fraction crosses the 0.5 threshold (Fig. 4E). These results provide evidence for a more general interpolation principle: regulatory conflicts are linearly interpolated for additive drug pairs, while antagonistic drug pairs elicit switch-like transitions, extending earlier transcription-level findings^36,37^ to the proteome.

Only a handful of proteins deviated qualitatively from the interpolation principle. This principle also does not accurately capture ribosome levels along isoboles (Fig. S9); instead, they align with the generalized growth law (Fig. 5). This is likely due to their exceptional regulatory mechanisms, which are more elaborate than those of most other proteins^7,49^. We found no consistent evidence of emergent responses, i.e., protein levels under the combination that exceeded or fell below both single-drug treatments. Any hints of such emergent responses under dual treatment (Fig. S8D) proved inconsistent across the dataset. Therefore, the proteome response to antibiotic combinations appears to be remarkably simple, as it is well captured by interpolating between the protein-level responses to the individual drugs. In practice, two ingredients are sufficient to predict the global proteome allocation under drug combinations: (i) the universal growth inhibition effect captured by the first PC, and (ii) each protein’s specific response to the individual antibiotics. Applying the interpolation principle together with a high-resolution growth-response surface enables the direct prediction of most combinatorial outcomes from the responses to individual drugs.

**Fig. 5:**
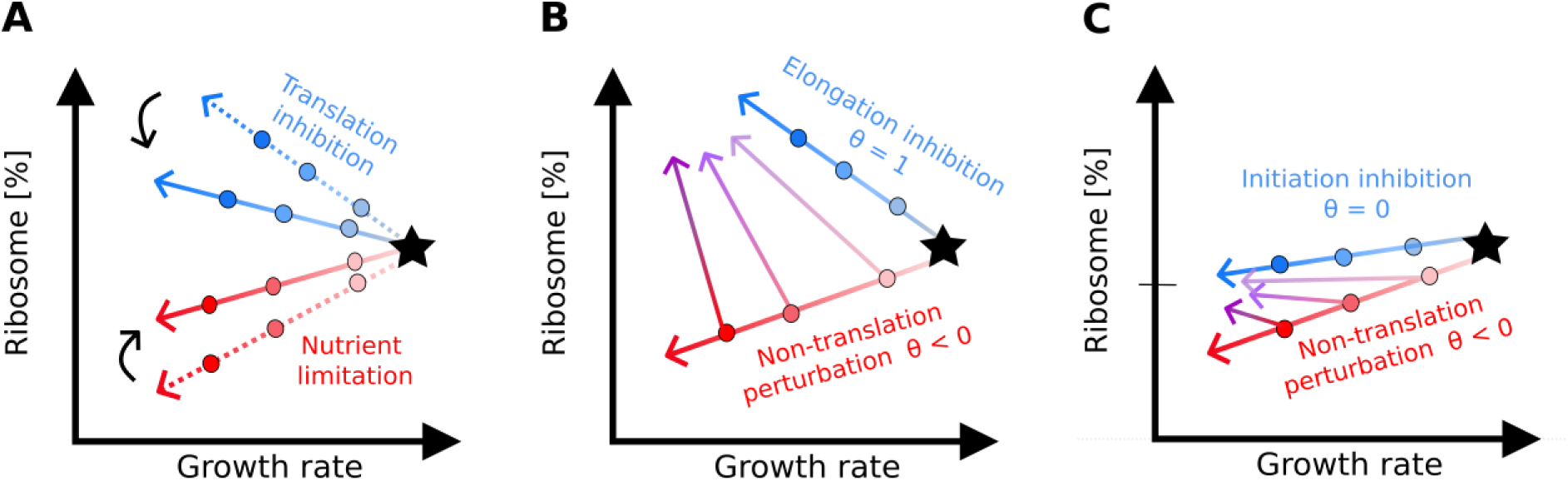
A generalized growth law unifies ribosome responses across combinations of perturbations. **A)** Schematic showing the first (dotted red arrow) and second bacterial growth law (dotted blue arrow). The solid lines illustrate the generalized growth law, in which the ribosome’s response to a non-translation perturbation (which does not directly target translation) and its response to translation inhibition can vary. **B,C)** Examples of specific predictions derived from the generalized growth law. Combining a non-translation perturbation with *θ* < 0 such as nutrient limitation or transcription inhibition (red) with a perturbation of translation elongation (*θ* = 1, blue) leads to ribosome responses similar to those of the second growth law, but with quantitative differences (B). Combining the non-translation perturbation with a perturbation of translation initiation or fidelity (*θ* ≈ 0) yields the ribosome responses depicted in C. These predictions generalize to perturbations with arbitrary values of *θ*.

## Discussion

Our results show that the bacterial proteome allocation response to translation-inhibiting antibiotics depends critically on the drug’s molecular mechanism of action and does not universally adhere to the second growth law. Inhibitors of translocation elongation or peptide bond formation produced the expected increase in ribosome concentration, whereas antibiotics targeting other stages of translation, such as initiation or decoding, resulted in a weaker or absent increase (Fig. 1B). In contrast, rifampicin-induced transcription inhibition triggered the opposite trend: ribosome levels declined while RNAP abundance increased (Fig. 1C). Under combined inhibition of transcription and translation, ribosome levels followed a superposition of the single-drug responses, closely resembling the second growth law under different nutrient conditions (Fig. 2). Incorporating this generalized growth law into a mathematical model of bacterial growth produced the graded levels of antagonism between rifampicin and different translation inhibitors (Fig. 3), indicating that drug interactions are at least partially shaped by stress-induced proteome reorganization. Building on earlier results at the transcriptional level^36,37,48^, we further showed that protein levels under antibiotic combinations are often smoothly interpolated along isoboles, rendering them partially predictable from the single-drug responses and growth-rate measurements alone (Fig. 4).

The divergent ribosome responses to distinct translation inhibitors likely arise from their different modes of action. In particular, these responses may be explained by signaling through the alarmones guanosine tetraphosphate and pentaphosphate, collectively termed (p)ppGpp, which represses ribosome production^50–52^. Synthesis of (p)ppGpp is stimulated under nutrient limitation, especially amino acid starvation, as uncharged tRNAs accumulate and activate the ribosome-associated (p)ppGpp synthetase RelA^51^. Inhibitors of translocation elongation or peptide bond formation, such as chloramphenicol, erythromycin, tetracycline, and doxycycline cause ribosomes to stall, thus reducing the fraction of active ribosomes^41^. The resulting surplus of charged tRNAs may suppress (p)ppGpp levels, thereby triggering the compensatory upregulation of ribosome production (Fig. 1B). This view aligns with the recent observation that (p)ppGpp levels are inversely related to the translation elongation rate across diverse growth conditions, suggesting that (p)ppGpp synthesis reflects the balance between dwelling ribosomes awaiting charged tRNAs and those undergoing productive translocation during elongation^29^. In contrast, drugs that block initiation, such as kasugamycin, or corrupt decoding fidelity, such as streptomycin, may not generate the stalled-ribosome signal needed to trigger compensatory ribosome production. To dissect these non-canonical responses (Fig. 1B), it will be essential to directly probe and perturb (p)ppGpp dynamics and other regulators of ribosome production^53^.

The decrease in ribosome concentration observed under rifampicin (Fig. 1B) is also mechanistically plausible. Rifampicin directly inhibits bacterial RNAP, reducing global mRNA synthesis and limiting the productive translation of all proteins, including ribosomal proteins^32^. Because ribosome biogenesis requires an exceptionally high level of transcription from rRNA operons^7,49^, a loss of transcriptional capacity disproportionately impairs rRNA synthesis. Since the rate of ribosome production is largely determined by rRNA synthesis^7,29,41^, rifampicin thereby limits the formation of new ribosomes, which offers a plausible explanation for the observed decrease in ribosome mass fraction relative to the proteome. This response is also consistent with recent theory proposing that growth under transcriptional limitation depends on both ribosome abundance and mRNA availability^30^. This work predicted a decrease in ribosome concentration in response to transcriptional inhibition in richer nutrient environments, which turns into an increase in poorer nutrient environments^30^. Our experiments used a defined medium with glucose as the carbon source (Methods), which supports a relatively high growth rate in *E. coli*, rendering the observed ribosome response broadly consistent with this prediction. An interesting direction for future work is testing the predicted effect of transcription inhibition by extending these quantitative proteomics measurements to poorer nutrient conditions.

Our results point to quantitative principles that govern bacterial responses to combinatorial perturbations in balanced exponential growth. First, the ribosome response to combinations of antibiotics that elicit distinct responses individually (Fig. 2) suggests that the second growth law may be a special case of a more general law. A non-translation perturbation – such as reduced nutrient quality or transcription inhibition – elicits a decline in ribosome abundance that depends linearly on growth rate (Fig. 5A). When translation is inhibited while holding the non-translation perturbation fixed, ribosome abundance again changes linearly with growth rate (Fig. 5B). Quantitatively, this second change mirrors the translation-inhibition response observed without the primary perturbation, but with a rescaled growth rate axis. Inhibitors of translocation elongation or peptide bond formation, such as chloramphenicol, elicit an increase in ribosome abundance, as predicted by the classical second growth law. However, the generalized growth law also captures the response to initiation inhibitors such as kasugamycin, and to other antibiotics for which ribosome abundance remains nearly constant. It makes concrete predictions, e.g., for the ribosome response to kasugamycin under different nutrient conditions (Fig. 5C). Whether this organizing principle applies to antibiotics with different modes of action or other stressors is an interesting question for future study.

Second, we found an interpolation principle that explains much of the proteome response to drug combinations based on the responses to the individual drugs (Fig. 4). This finding builds upon earlier studies in *E. coli* and yeast showing that transcriptional responses to antibiotic combinations occupy a low-dimensional space largely determined by the single-drug responses^36,37,48^. We found that the same principle extends to the proteome, the more functionally relevant layer. After factoring out the dominant growth-rate effect, additive drug pairs exhibit near-linear interpolation between single-drug proteome states, whereas antagonistic combinations produce more switch-like transitions between distinct physiological programs (Fig. 4B,D). The first PC (growth rate) explained a smaller fraction of the variance than in prior transcriptomic studies (Fig. 4A), likely because protein levels are subject to tighter post-transcriptional regulation, and proteomic sampling is sparser and noisier. Denser sampling along growth isoboles^37^ could sharpen the extraction of low-dimensional proteome response modes. Collectively, these findings indicate that large swaths of the bacterial response to antibiotic combinations can be predicted from single-drug data, offering a strategy to avoid the combinatorial explosion in drug-combination design.

Although our study examined several mechanistically distinct antibiotics, the broader landscape of translation and transcription inhibitors offers fertile ground for future work. Several inhibitors act through distinct mechanisms that may help further elucidate the interplay between translation and transcription in bacteria. For example, we did not investigate transcription elongation inhibitors such as streptolydigin, which perturbs RNAP through a mechanism that differs from that of rifampicin^32,54,55^. Similarly, aminoacyl-tRNA synthetase inhibitors such as mupirocin, which strongly activate the stringent response through accumulation of uncharged tRNAs^50–52,56^, could elicit markedly different proteome allocation states. The same applies to translation inhibitors that cause premature chain termination and proteotoxic stress, such as puromycin^57^. Applying quantitative proteomics to these antibiotics and their combinations would further challenge the universality of the identified principles of bacterial physiology.

The fact that biophysical models of resource allocation extend so readily from translation to transcription inhibition underscores the power of empirical growth laws derived from quantitative proteomics. Further advances in proteomics methodology, such as improved coverage, more accurate absolute quantification, and robust cross-species normalization, could extend this framework to antibiotics targeting other cellular processes, and to bacteria beyond *E. coli*. In other species, the relationship between ribosome abundance and growth rate may differ^10^, and regulators other than (p)ppGpp may take center stage^58^. Combining these advances with high-throughput, automated sample preparation will enable rigorous testing of whether system-level antibiotic stress responses are universally governed by low-dimensional physiological constraints. Ultimately, this approach could transform the design of effective antibiotic combination therapies from a process of trial-and-error to one of quantitative prediction.

## Methods

### Bacterial strains

Experiments were performed using *Escherichia coli* K-12 BW25113 carrying the pCS-λ plasmid, which encodes the constitutive *luxCDABE* reporter system. Calibration measurements were performed using the EQ353 strain, as described in^14,39^.

### Growth media

Unless otherwise stated, bacteria were grown in M9 minimal medium consisting of 1×M9 salts (Sigma-Aldrich, M6030), supplemented with 2 mM MgSO₄ (Sigma-Aldrich, M7506), 0.1 mM CaCl_2_ (VWR, 22317), and 4 g/L glucose (Sigma-Aldrich, G8270). Carbon limitation experiments were performed by increasing the concentration of methyl α-D-glucopyranoside (Sigma-Aldrich, M9376-1KG) while keeping the glucose concentration constant. This resulted in methyl α-D-glucoside/glucose ratios of 5, 10, 15, and 20. For the calibration experiments, cultures of the EQ353 strain were grown in MOPS minimal medium prepared as described in^39^, using MOPS (Sigma-Aldrich, M3183-500G).

### Antibiotics

We used rifampicin (Sigma-Aldrich, R3501-1G), chloramphenicol (Sigma-Aldrich, C0378), kanamycin (Sigma-Aldrich, K4000), streptomycin sulfate (Sigma-Aldrich, S6501-25G), tetracycline (Sigma-Aldrich, 87128), doxycycline hyclate (Sigma-Aldrich, D9891-1G), kasugamycin hydrochloride from *Streptomyces kasugaensis* (Sigma-Aldrich, K4013-10G), and erythromycin hydrate (Sigma-Aldrich, 856193-25G). Stock solutions of rifampicin, streptomycin sulfate, and kasugamycin were prepared in pure water (MilliQ).

Chloramphenicol, tetracycline, doxycycline, and erythromycin stock solutions were prepared in ethanol. All stock solutions were filter sterilized and stored at −20°C.

### Bioreactor and batch culture growth assays for proteomics samples

#### Batch culture experiments

Batch culture experiments were performed in 100 mL Erlenmeyer flasks containing 20 mL of medium. The flasks were incubated in a shaking incubator at 37°C and 250 rpm. Cultures were inoculated from frozen stocks by 1:1,000 dilution into a pre-culture medium that was identical to the respective experimental conditions. During the exponential growth phase, the cultures were further diluted 1:100 into fresh medium to obtain the desired starting conditions. Growth was monitored by measuring the optical density at 600 nm (OD₆₀₀) using a spectrophotometer (NanoDrop OneC). Samples were harvested for further protein extraction at OD_600_ = 0.20.

#### Bioreactor experiments

Aliquots of frozen overnight cultures (50 µL in 15% glycerol) were thawed and inoculated at a 1:10,000 dilution into flasks containing 15 mL of growth medium connected to the mini-bioreactor system (OGI Biotec, Bioreactor Mk3 Main CCA). Growth was continuously monitored. Samples (10 mL) for proteomics analysis were collected at OD₆₀₀ = 0.2 to ensure sampling during the exponential growth phase. The bioreactor system estimates cell density using a backscatter signal, which was calibrated against spectrophotometric OD₆₀₀ measurements using a power-law relationship 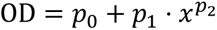, where (*x*) denotes the measured backscatter signal. All bioreactor experiments were performed at 37°C.

### Proteomics sample processing

Samples were centrifuged for 12 min at 3,000 × *g* at 4°C using a refrigerated centrifuge. The supernatant was discarded, and cell pellets were washed with 2 mL PBS buffer (Carl Roth, 1107.1). The pellets were then centrifuged again for 2 min at 13,000 × *g* using a tabletop centrifuge and subsequently stored at −80°C until protein extraction.

#### Protein quantification

Total protein concentration was determined using a modified Lowry assay (Total Protein Kit Micro Lowry, Peterson’s modification; Sigma-Aldrich, TP0300-1KT), which was adapted for small sample volumes.

#### Proteomics sample preparation

We prepared proteomics samples according to the standard protocol of the CECAD/CMMC Proteomics Facility at the University of Cologne. Cell pellets were washed with ice-cold PBS and resuspended in 100 µL lysis buffer containing 100 mM ammonium bicarbonate (Sigma-Aldrich, A6141), 2% sodium deoxycholate (Sigma-Aldrich, D6750), and protease inhibitor cocktail (cOmplete, Roche, 04693159001). Cells were lysed by vortexing and sonication on ice. Cell debris was then removed by centrifugation (16,000 × g, 15 min, 4 °C). The supernatant was transferred to a fresh tube and stored at −80 °C until further processing. Protein concentration was determined prior to digestion. Approximately 30 µg protein was reduced with 5 mM dithiothreitol (DTT) for 1 h at 20 °C and alkylated with 40 mM chloroacetamide (CAA) for 30 min at 30°C in the dark. Proteins were digested with Lys-C at an enzyme-to-substrate ratio of 1:75 for 4 h at 25°C, followed by overnight digestion with trypsin at a ratio of 1:75 at 25°C. Digestion was stopped by adding trifluoroacetic acid to a final concentration of 5%, followed by centrifugation to remove precipitated detergent. Peptides were purified using SDR StageTips preconditioned with methanol and equilibrated with buffer A (0.1% formic acid in water) and buffer B (0.1% formic acid in 80% acetonitrile). Samples were washed and eluted according to the facility protocol, then stored at 4 °C until LC–MS analysis.

#### Data Acquisition

Samples were analyzed by the CECAD Proteomics Facility on an Orbitrap Exploris 480 mass spectrometer either coupled to a Vanquish neo in reverse flow setup (all Thermo Scientific) or an Evosep One (Evosep). For analyses using a Vanquish neo, samples were loaded onto a PepMap precolumn cartridge (#160434, Thermo Scientific) before reverse-flushing onto an in-house packed 30 cm column filled with C18 material (75 µm inner diameter, filled with Agilent Poroshell). Samples were separated on a 90 min gradient running eluent A (0.1% formic acid) against eluent B (80% acetonitrile + 0.1% formic acid) on a linear gradient between 4% and 98% B. Alternatively, samples were loaded onto Evotips and analyzed using the 30 SPD method with vendor-recommended column on the Evosep One. The mass spectrometer was operated in data-independent acquisition mode with MS1 scans acquired from 399 m/z to 1001 m/z, MS2 scans ranged from 400 m/z to 1000 m/z. Both MS1 and MS2 scans were acquired at 15k or 30k resolution, respectively, with maximum injection times set to allow maximal parallelization. AGC targets for MS2 were set to 1000% and MS2 scans were acquired either in 60 x 10 m/z or 30 x 20 m/z windows with an overlap of 1 m/z. All scans were stored as centroid.

For the Gas-phase fractionated library^59^, a pool generated from all samples was analyzed in six individual runs covering the range from 400 m/z to 1000 m/z in 100 m/z increments using identical LC settings as the samples. For each run, MS1 was acquired at 60k resolution with a maximum injection time of 98 ms and an AGC target of 100%. MS2 spectra were acquired at 30k resolution with a maximum injection time of 60 ms. Spectra were acquired in staggered 4 m/z windows, resulting in nominal 2 m/z windows after deconvolution using ProteoWizard^60^.

#### Sample Processing in DIA-NN

The gas-phase fractionated library was built using DIA-NN 1.8.1^61^ using a Swissprot *E. coli* canonical database (downloaded 04/01/2023 and 15/01/2025, respectively) with settings matching acquisition parameters. DIA-NN was run with the additional command line prompts “—report-lib-info” and “—relaxed-prot-inf”. Further output settings were: filtered at 0.01 FDR, N-terminal methionine excision enabled, maximum number of missed cleavages set to 1, min peptide length set to 7, max peptide length set to 30, min precursor m/z set to 400, max precursor m/z set to 1000, cysteine carbamidomethylation enabled as a fixed modification. Finally, acquired sample files were searched against the GPF library or, alternatively, library-free using the match-between-runs option in DIA-NN. Afterwards, DIA-NN output was further filtered on library q-value and global q-value <= 0.01 and at least one unique peptides per protein using R (4.1.3). The mass spectrometry proteomics data have been deposited to the ProteomeXchange Consortium via the PRIDE partner repository^62^ with the dataset identifier PXD082778.

### Quantitative protein abundance analysis

Peptide abundances were aggregated into protein abundances using the xTop v2 algorithm as detailed in^14^. Protein molecular weights were obtained from UniProt and used to convert protein abundances into proteome mass fractions by multiplying each protein abundance by its corresponding molecular weight and normalizing by the total summed protein mass. To determine absolute proteome mass fractions, we included calibration conditions for which ribosome profiling reference data are available^39^ in the dataset. Protein abundances were scaled to these reference conditions following the calibration procedure described in^14^. Proteins were assigned to proteome sectors as previously described^40^. The flask dataset exhibited a global median coefficient of variation (CV) of approximately 14%, while replicated anchor conditions in the mini-bioreactor dataset exhibited a comparable median CV of approximately 16%. Samples with biological replicates were averaged, while conditions without replicates are represented by a single measurement. The number of replicates is indicated in the column headings of Tables S1 and S2.

### Global proteome analysis

Protein abundances for each drug combination condition were normalized to the corresponding untreated condition. Prior to downstream analysis, proteins containing missing values or zero abundances across analyzed conditions were excluded. Relative protein abundances were transformed into log_2_ fold changes relative to the untreated condition. For principal component analysis (PCA), protein abundance profiles were standardized using `sklearn.preprocessing.StandardScaler`. PCA was subsequently performed using `sklearn.decomposition.PCA` to identify dominant axes of proteome variation across drug-treatment conditions. To determine which functional groups contributed the most to each principal component, proteins were grouped by their level-2 annotation. For each group, squared PC scores were summed and normalized to the total squared scores of all level-2 annotated proteins. Positive and negative scores were considered separately to preserve the direction of the contribution. Drug concentrations were normalized by their IC_50_, determined from single-drug dose-response measurements using a Hill function fitted with `scipy.optimize.curve_fit`:

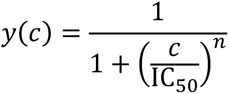

where *c* denotes drug concentration, IC_50_ the half-maximal inhibitory concentration, and *n* the Hill coefficient.

Continuous response surfaces for growth rate, principal component scores, and individual protein abundances were reconstructed on dense, regular grids using `numpy.meshgrid` and piecewise linear interpolation with `scipy.interpolate.griddata` (`method="linear"`). Interpolation was performed in IC_50_-normalized concentration space and restricted to the convex hull of experimentally measured conditions to avoid extrapolation beyond sampled concentration regions. Isoboles corresponding to constant growth inhibition levels were extracted from interpolated growth surfaces and plotted as a function of the relative drug fraction 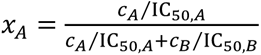 (Figs. 4B,D,E and Fig. S5F).

Protein trajectories along isoboles were analyzed using both linear and sigmoidal models. Linear trajectories were fitted using ordinary least-squares regression implemented with `scipy.stats.linregress`. The linear fits were then evaluated using the coefficient of determination (*R*^2^), slope directionality, and monotonicity along the analyzed trajectories. Sigmoidal trajectories were fitted using `scipy.optimize.curve_fit` and a hyperbolictangent function:

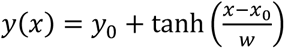

where *y₀* denotes the baseline level, *h* the transition amplitude, *x*₀ the midpoint position, and *w* the transition width parameter. Model quality was evaluated using the coefficient of determination (R^2^) and the Akaike Information Criterion (AIC). We compared AIC values between sigmoidal and linear fits to assess if the sigmoidal fits substantially improved the description of the data. Proteins were classified only if at least five valid data points were available along the analyzed isobole and if the dynamic range exceeded 0.30 log_2_ fold-change units. Proteins were classified as sigmoidal if they satisfied all of the following criteria: *R*^2^ ≥ 0.90 for the sigmoidal fit, ΔAIC ≥ 4 compared to the linear model, transition width *w* < 0.40, and midpoint position 0.45 ≤ *x*₀ ≤ 0.55. Proteins were classified as linear if they satisfied the following criteria: linear fit quality *R*^2^ ≥ 0.90 and monotonic fraction ≥ 0.6. The monotonic fraction was defined as the fraction of neighboring steps along the isobole that followed the dominant direction of change. Proteins not fulfilling these criteria were classified as unassigned. To reduce computational costs during large-scale fitting, proteins exhibiting highly linear behavior (linear *R*^2^ > 0.97) were optionally excluded from additional sigmoidal fitting steps.

#### Candidate emergent-response examples

To identify possible emergent responses, protein trajectories along the analyzed isobole were screened for proteins with higher abundance in the mixed-drug region than at both single-drug ends. The single-drug edge regions were defined as relative drug fractions x≤0.15 and x≥0.85, and the mixed-drug region as 0.35≤*x*≤0.65. Candidate proteins were required to have at least five valid data points, edge values close to baseline, and a higher maximum log_2_ fold-change in the mixed-drug region than at either edge. The four clearest candidates were selected for visualization in Fig. S5D, but because this behavior was weak and inconsistent across the dataset, they were treated as illustrative examples rather than as a robust emergent-response class.

### Quantification of drug interactions

Drug interactions were quantified using the logarithmic Loewe interaction score, LI_log_, following the volume-based definition of Kavčič *et al.*^2^. For each antibiotic pair, the Loewe-additive response surface, *y*_add_(*c_A_*, *c_B_*), was reconstructed from the normalized single-drug dose-response curves. At each concentration pair, the additive growth response *y*(*c_A_*, *c_B_*) was determined by solving

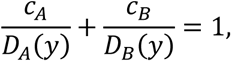

where *D_i_*(*y*) is the concentration of drug *i* required to produce the normalized growth response *y*. Within each analysis, the measured or model-predicted response surface and its corresponding additive surface were integrated over identical points in two-dimensional concentration space. The interaction score was calculated as

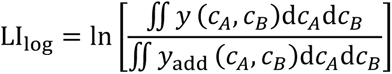

Experimental surfaces were integrated over the original measured concentration grid using weights determined by the spacing between concentrations, with missing values excluded. Model surfaces were evaluated on an 80×80 grid spanning 0–1.6×IC_50_ along both drug axes. Positive LI_log_ values indicate antagonism, negative values indicate synergy, and values close to zero indicate additivity.

### Numerical solutions

We evaluated the steady-state solutions of the one- and two-drug models described in the Supplementary Text by directly solving the corresponding cubic equation for the growth rate λ. All roots were calculated using `numpy.roots`, and only real roots within the physically admissible range 0≤λ≤λ_0_ were retained. At zero drug concentration, the largest admissible root was selected. At subsequent concentrations, the root closest to the preceding solution was chosen to follow a continuous solution branch. Growth was normalized by the drug-free growth rate, *y*=λ/λ_00_, and the model IC_50_ was determined by interpolation at *y*=0.5.

For the two-drug model, the transcription inhibitor X established a concentration-dependent background growth rate, *λ*_0_(*c_X_*) = *λ*_00_*g_X_*(*c_X_*), described by a normalized Hill function with IC_50,X_=0.7 and Hill coefficient *n*_X_=2. At each concentration of drug X, the effective ribosome range was recalculated according to the generalized growth law, and the steady-state equation was solved over increasing concentrations of the translation inhibitor Y. We used *κ*_t_=0.061μM^−1^h^−1^, *r*_min_=19.3μM, Δ*r*=46.5μM, and λ_00_=0.85h^−1^; the remaining kinetic parameters were set as described in the Supplementary Text. Two-dimensional response surfaces were calculated for *θ*=1 and *θ*=0, representing strong and weak ribosome upregulation, respectively. For Fig. S4A, one-dimensional dose-response curves were calculated for *θ*=−0.4, 0, and 1, with concentrations normalized by the IC_50_ of the *θ*=0 response.

## Data Availability

Source data are provided with this paper. Proteomics raw data files have been uploaded to ProteomeXchange under the study accession code PXD082778. Proteome sectors and slopes of conditions are provided in the Supplementary Information (Table S1). Also provided in the Supplementary Information are the expression levels of all detected proteins in all conditions investigated here (Table S2).

## Code Availability

This study did not produce any new computer code, except for a few Python 3 scripts used for data analysis and exploration. All codes are available from the corresponding author upon request.

## Acknowledgments

We thank the entire Bollenbach group, especially Booshini Fernando, for technical support and fruitful discussions; Marco Cosentino-Lagomarsino, Theresa Fink, Terence Hwa, and Leon Seeger for critical reading of the manuscript; and Jan-Wilm Lackmann and the whole CECAD / CMMC Proteomics Facility for technical support.

## Author contributions

Conceptualization: N.G. and T.B. Investigation: N.G. Formal Analysis: N.G. and M.M. Writing – original draft: N.G. and T.B. Writing – review and editing: N.G., M.M., and T.B. Supervision: T.B. Funding acquisition: T.B.

## Supporting Information

### Supplementary figures

**Fig. S1:**
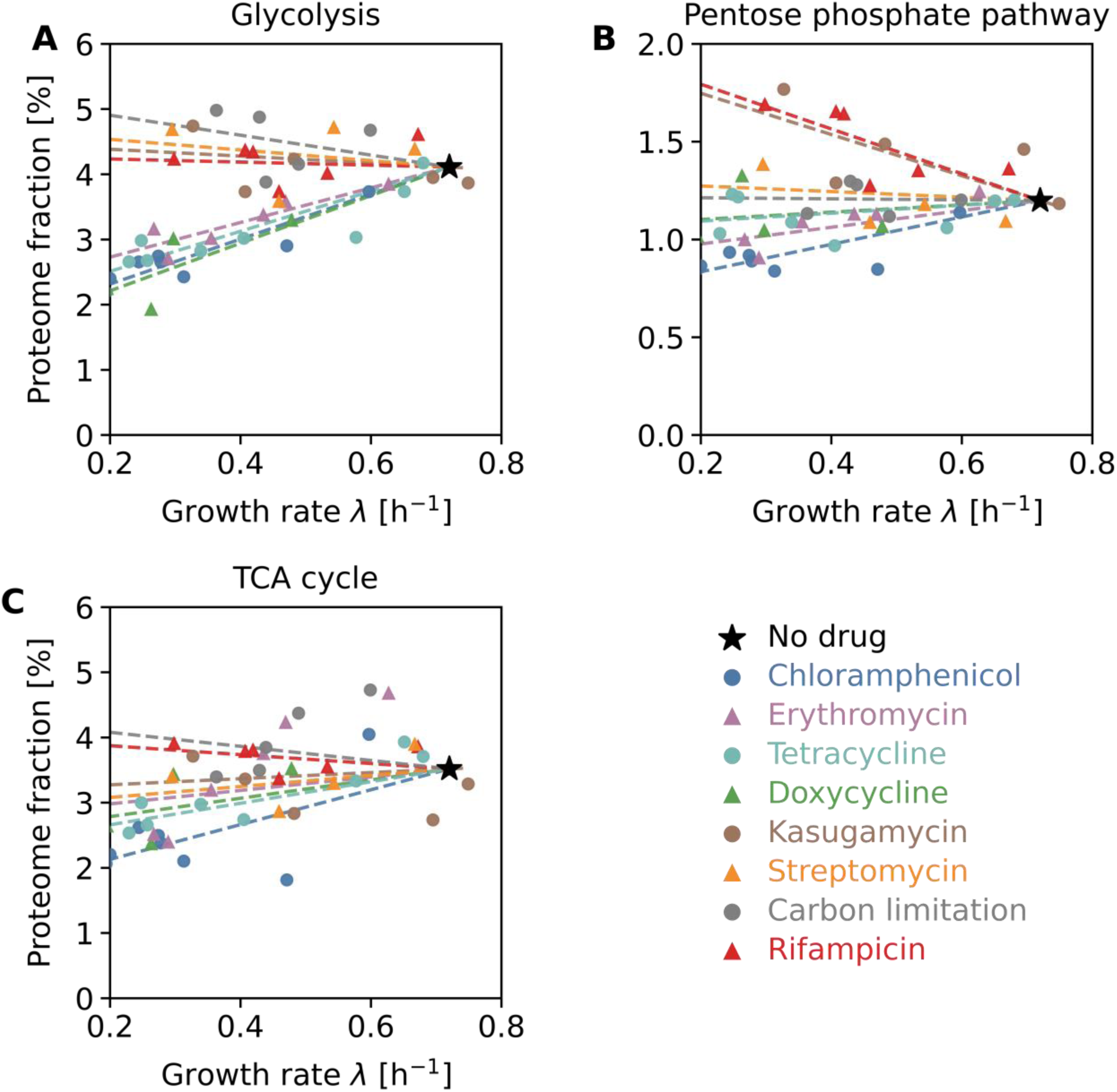
Drug-specific sector responses of central carbon metabolism. **A)** Glycolysis protein mass fraction as a function of growth rate under chloramphenicol (CHL), kasugamycin (KSG), rifampicin (RIF), and carbon limitation. **B)** As in A, but for proteins belonging to the pentose phosphate pathway. **C)** As in A and B, but for proteins belonging to TCA cycle. Data were obtained from bioreactor-grown cultures (Methods).

**Fig. S2:**
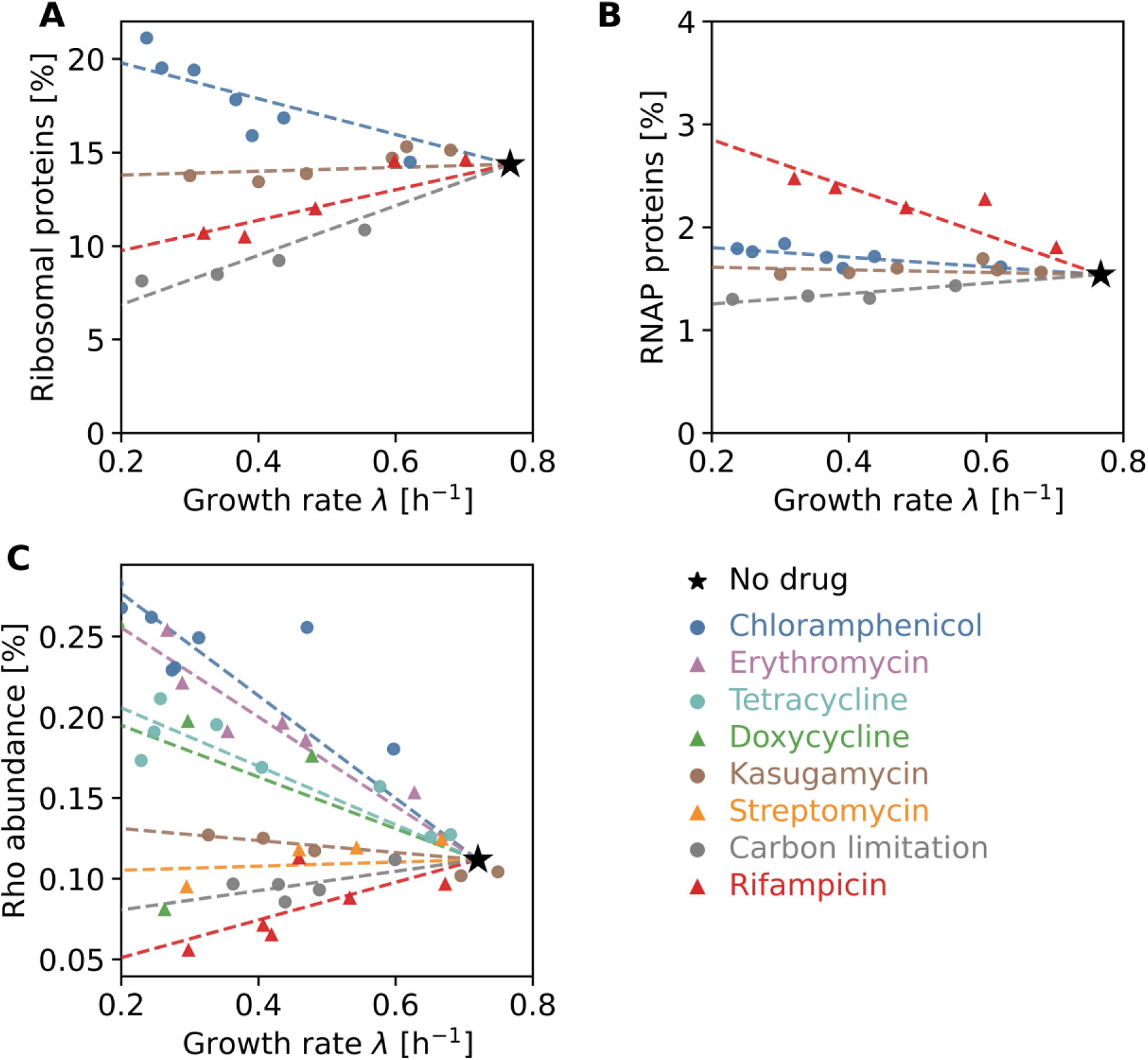
Independent replicate proteomics dataset for one-dimensional gradients of antibiotics and carbon limitation. **A)** Ribosomal protein mass fraction as a function of growth rate under chloramphenicol (CHL), kasugamycin (KSG), rifampicin (RIF), and carbon limitation, as in Fig. 1B, but for an independent dataset. **B)** RNAP-associated protein mass fraction as in Fig. 1C, but for an independent dataset. Rifampicin increases RNAP allocation, whereas translation inhibitors produce comparatively minor changes. **C)** As Fig. 1B,C, but for the transcription-termination factor Rho. Rho follows a similar trend to that of ribosomal proteins across all perturbations. Data in A and B were obtained from cultures grown in flasks, whereas data in C were obtained from bioreactor-grown cultures (Methods).

**Fig. S3:**
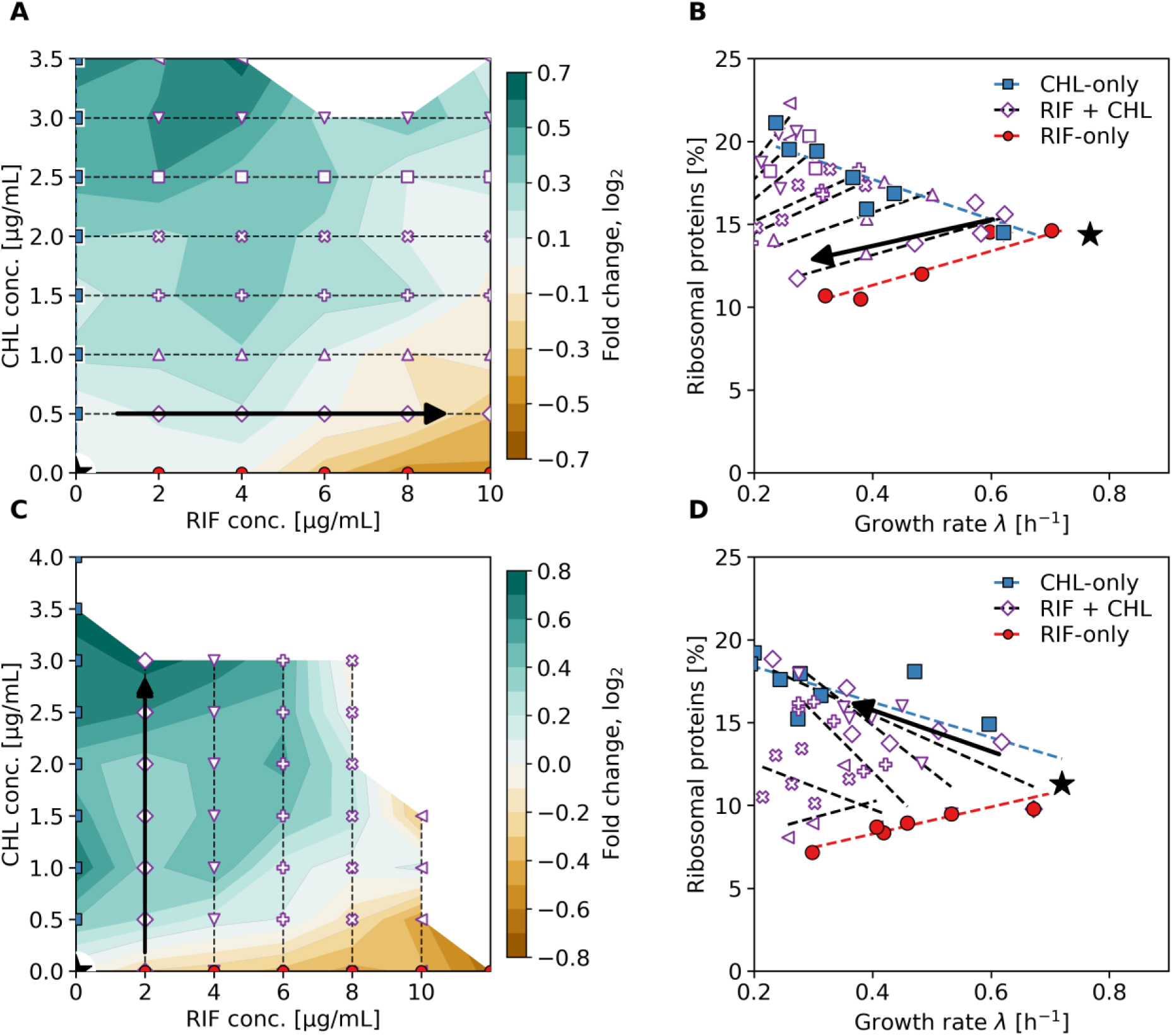
Ribosome response to transcription inhibition at fixed translation inhibition and independent replicate proteomics measurement of the response to rifampicin-chloramphenicol. **A)** Heatmap of ribosomal protein abundance (fold change relative to the no-drug condition) across a two-dimensional concentration gradient of rifampicin (RIF) and chloramphenicol (CHL). Black arrow indicates trajectory of increasing rifampicin concentration at a fixed chloramphenicol concentration. Ribosome allocation continuously shifts from the elevated ribosome state characteristic of chloramphenicol treatment toward the low-ribosome rifampicin state. **B)** Ribosomal protein mass fraction as a function of growth rate for rifampicin-only, chloramphenicol-only, and combined rifampicin-chloramphenicol conditions at the concentrations indicated in A. **C)** As Fig. 2A, but measured in an independent steady-state bioreactor dataset. Black arrow indicates trajectory of increasing chloramphenicol concentration at a fixed RIF concentration. **D)** As Fig. 2B, but for the independent dataset in C. Combined-drug conditions interpolate between the single-drug growth-law responses, reproducing the trends observed in Fig. 2. Data in A and B were obtained from cultures grown in flasks, whereas data in C and D were obtained from bioreactor-grown cultures (Methods).

**Fig. S4:**
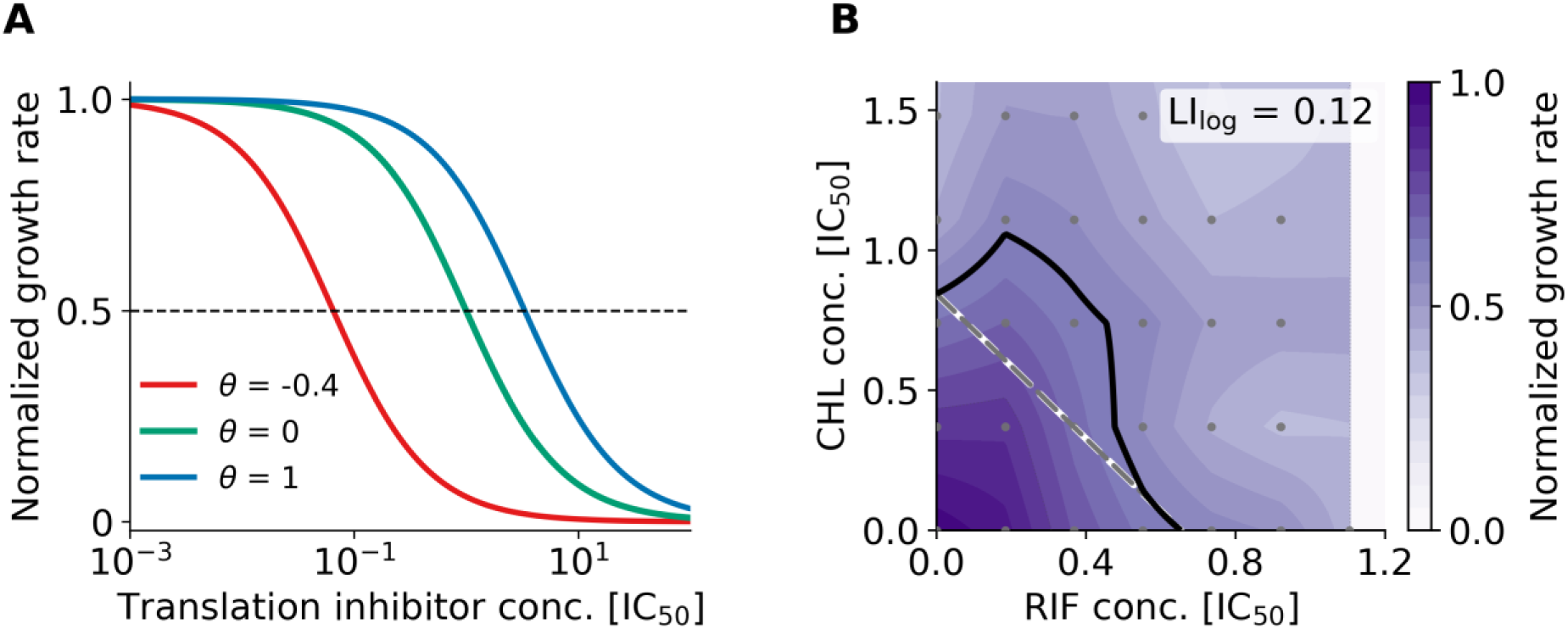
Properties and independent experimental validation of the mathematical model based on the generalized growth law. **A)** Dose-response curves calculated from the mathematical model for translation inhibitors with different ribosome-response parameters *θ*. Changing the value of *θ* rescales the effective drug concentration and shifts the dose-response curve horizontally without substantially altering its shape. Strong ribosome upregulation (*θ*=1) decreases susceptibility, whereas weak or negative ribosome responses increase susceptibility. **B)** Independent bioreactor replicate of the rifampicin (RIF) and chloramphenicol (CHL) growth-response surface (Methods). The antagonistic interaction observed in Fig. 3B is reproduced. Dashed line: Loewe-additive expectation. Gray points indicate experimentally measured conditions. Data in B were obtained from bioreactor-grown-cultures (Methods).

**Fig. S5:**
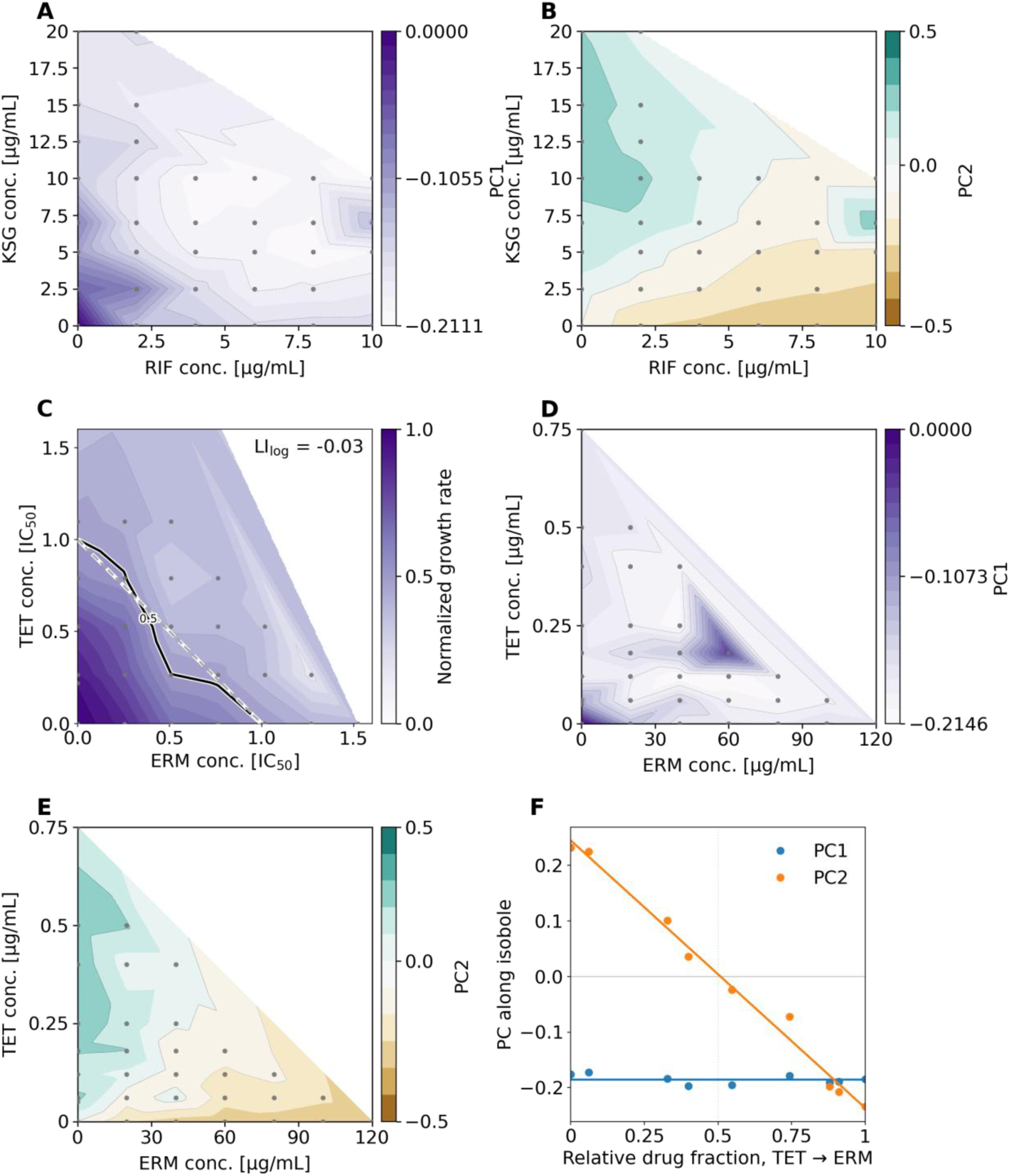
Growth response surfaces and principal components of proteome responses for additive combinations of translation and transcription inhibitors. **A-B)** First (A) and second (B) principal component for the rifampicin-kasugamycin (RIF-KSG) combination. **C-D)** Growth-response surface (C) and first principal component (D) for the erythromycin-tetracycline (ERM-TET) combination. **E-F)** Second principal component in two-dimensional concentration space (E) and along the isobole at 50% growth inhibition (F) for the erythromycin-tetracyclline combination. Data in A and B were obtained from cultures grown in flasks, whereas data in C–F were obtained from bioreactor-grown cultures (Methods).

**Fig. S6:**
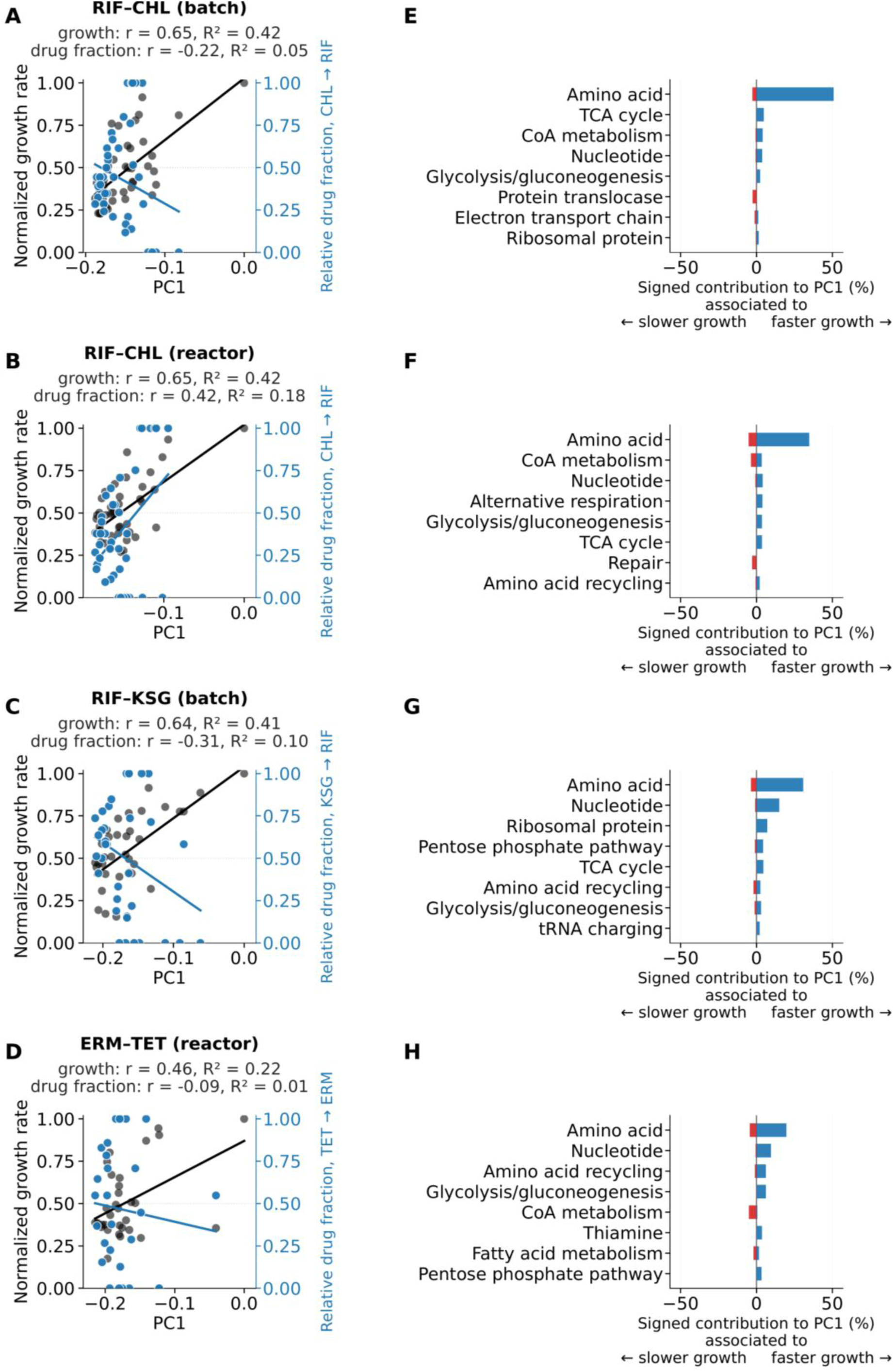
The first principal component captures growth-rate-dependent proteome variation. **A–D)** normalized growth rate (gray) and relative drug fraction (blue) plotted against the first PC (PC1) for rifampicin–chloramphenicol (A, batch culture; B, bioreactor), rifampicin– kasugamycin (C, batch culture), and erythromycin–tetracycline (D, bioreactor). Lines show linear fits; Pearson’s *r* and *R*^2^ are indicated. The relative fraction of drug A in a combination with drug B is defined as 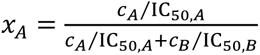 (Methods). **E–H)** Signed contributions to the first PC of the main level-2 functional groups for rifampicin–chloramphenicol (E, batch culture; F, bioreactor), rifampicin–kasugamycin (G, batch culture), and erythromycin–tetracycline (H, bioreactor). Positive values are associated with faster growth and negative values with slower growth.

**Fig. S7:**
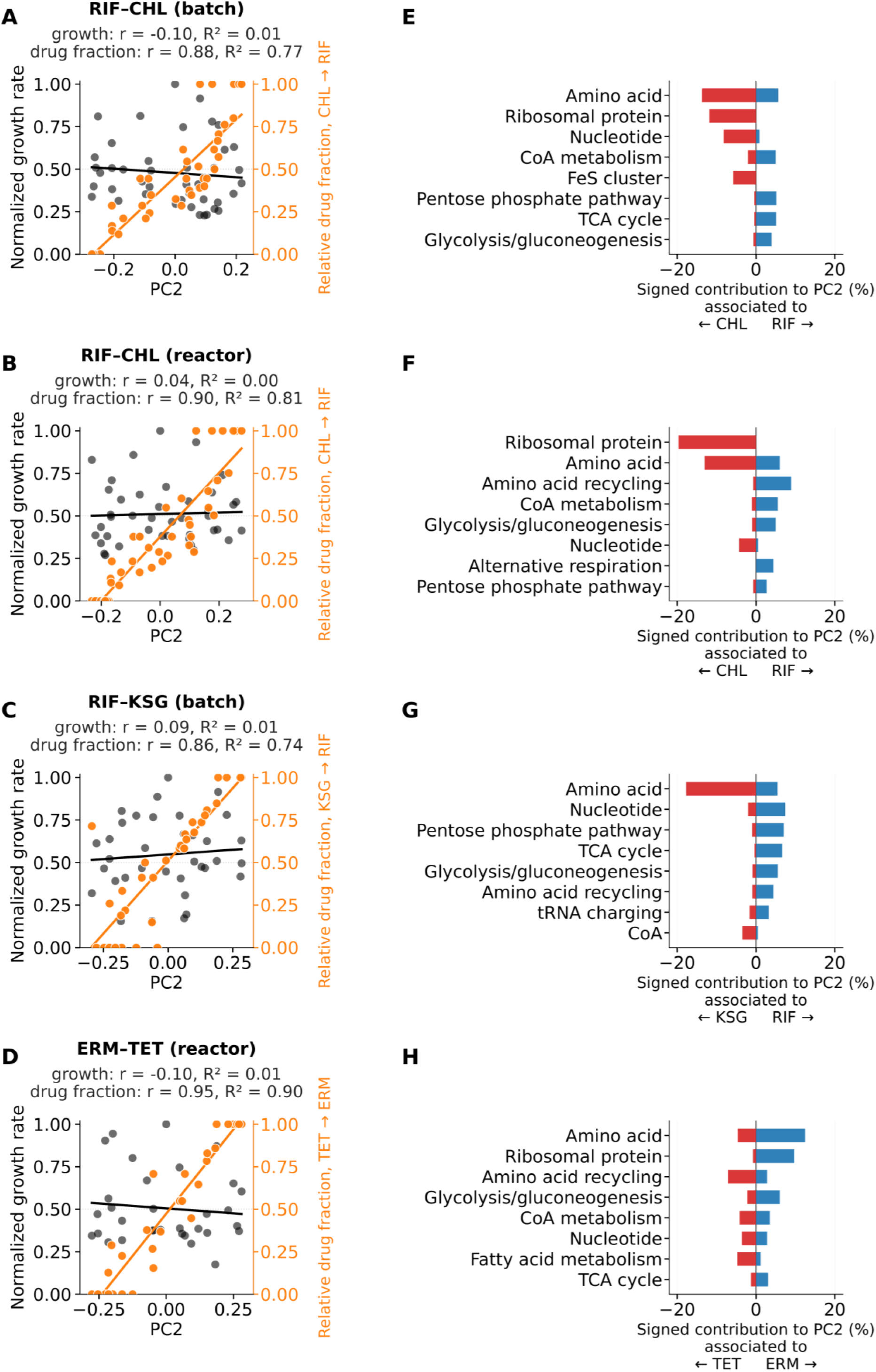
The second principal component captures drug-specific proteome variation. **A–D)** Second PC plotted against normalized growth rate (gray) and relative drug fraction (orange) for the same four datasets as in Fig. S6A-D. Lines show linear fits; Pearson *r* and *R*^2^ are indicated. **E–H)** Signed contributions to the second PC of the main level-2 functional groups. The direction of the contribution indicates association with the drug indicated below each panel.

**Fig. S8:**
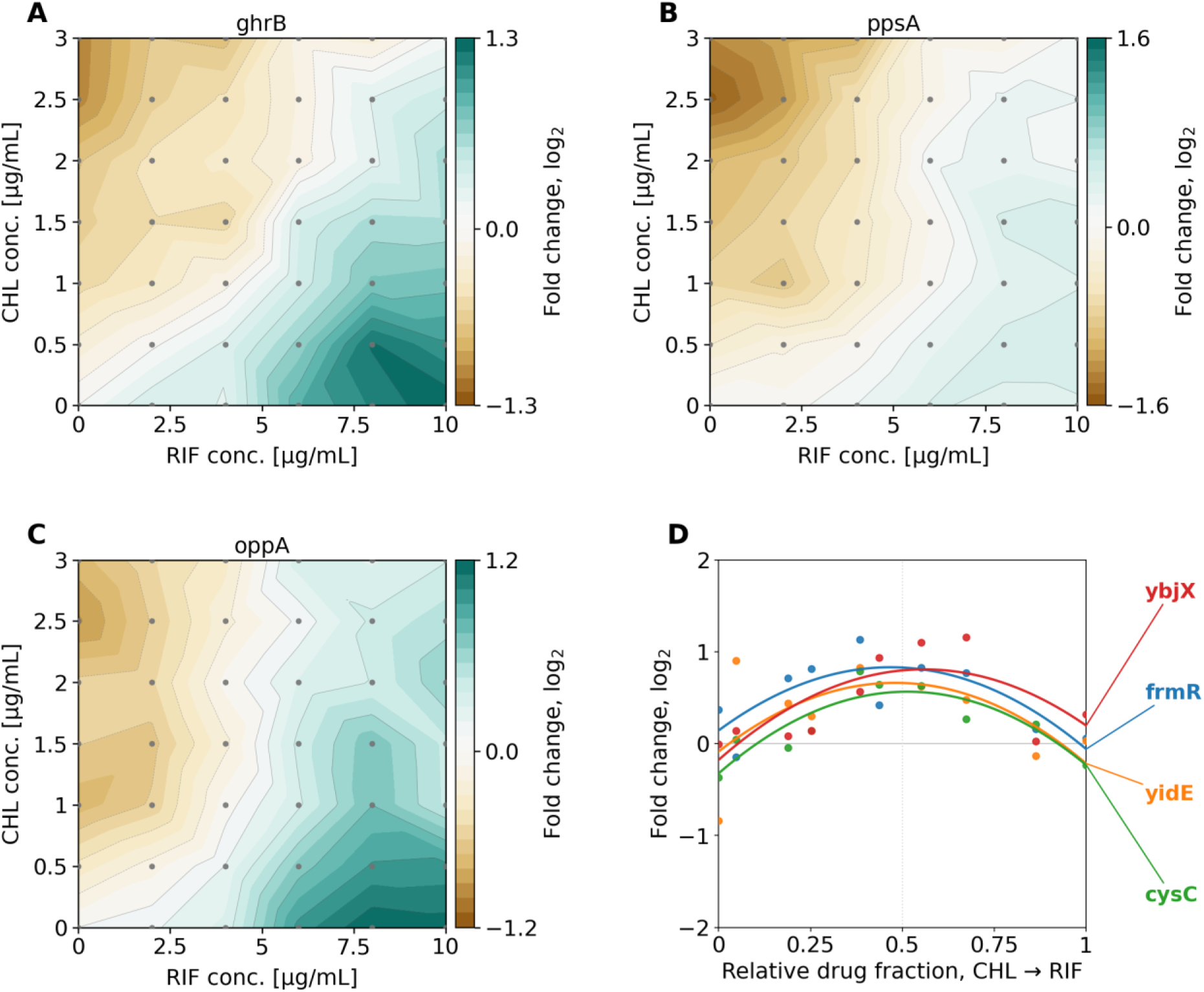
Examples of protein-level responses to an antibiotic combination and candidates for emergent responses. **A-C)** Fold-change in expression level in two-dimensional concentration gradient of chloramphenicol (CHL) and rifampicin (RIF) for the glyoxylate reductase GhrB (A), the phosphoenolpyruvate synthetase PpsA (B), and OppA (C), the periplasmic binding protein of an oligopeptide ABC transporter^49^. **D)** Expression level along isobole as in Fig. 4E for the four best candidate proteins (YbjX, FrmR, YidE, CysC) exhibiting signs of emergent responses (Methods). Data in A–D were obtained from bioreactor-grown cultures (Methods).

**Fig. S9:**
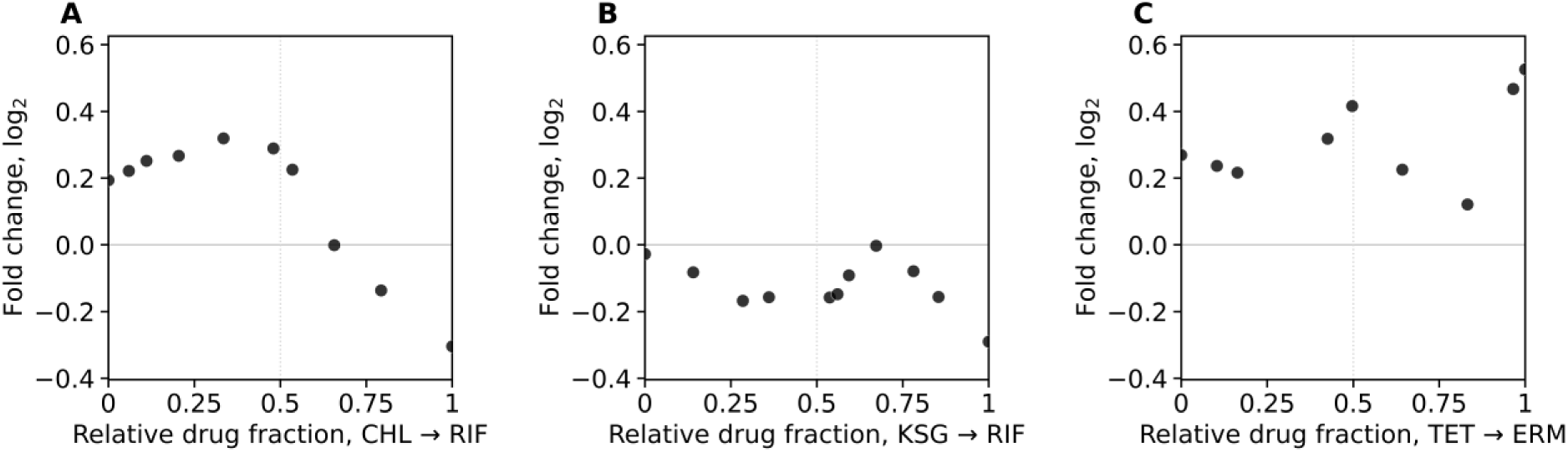
Ribosome response to antibiotic combinations along isoboles. **A)** Fold-change in ribosome concentration in response to the rifampicin-chloramphenicol (RIF-CHL) combination as in Fig 2A, but plotted along the isobole at 50% growth inhibition as in Fig. 4E. **B-C)** As A, but for the rifampicin-kasugamycin (RIF-KSG, B) and erythromycin-tetracycline (ERM-TET) combinations (C). Data in A and B were obtained from cultures grown in flasks, whereas data in C were obtained from bioreactor-grown cultures (Methods).

### Supplementary data

**Table S1:** Assignments of proteins and slopes in one-dimensional antibiotic concentration gradients for different sectors in level 1–3 groups across two independent batches^40^.

**Table S2:** Absolute scaled protein mass fractions (in percent) for all detected proteins across all experimental conditions.

## References

1. Greulich, P., Scott, M., Evans, M. R. & Allen, R. J. Growth-dependent bacterial susceptibility to ribosome-targeting antibiotics. Mol Syst Biol 11, MSB145949 (2015).

2. Kavčič, B., Tkačik, G. & Bollenbach, T. Mechanisms of drug interactions between translation-inhibiting antibiotics. Nat Commun 11, 4013 (2020).

3. Kavčič, B., Tkačik, G. & Bollenbach, T. Minimal biophysical model of combined antibiotic action. PLoS Comput Biol 17, e1008529 (2021).

4. Droghetti, R. et al. Hands-On Growth Laws Theory Cookbook. PRX Life 4, 012001 (2026).

5. Scott, M., Gunderson, C. W., Mateescu, E. M., Zhang, Z. & Hwa, T. Interdependence of Cell Growth and Gene Expression: Origins and Consequences. Science 330, 1099–1102 (2010).

6. Schaechter, M., Maaløe, O. & Kjeldgaard, N. O. Dependency on Medium and Temperature of Cell Size and Chemical Composition during Balanced Growth of Salmonella typhimurium. Journal of General Microbiology 19, 592–606 (1958).

7. Bremer, H. & Dennis, P. P. Modulation of Chemical Composition and Other Parameters of the Cell at Different Exponential Growth Rates. EcoSal Plus 3, 10.1128/ecosal.5.2.3 (2008).

8. Dennis, P. P. & Bremer, H. Macromolecular Composition During Steady-State Growth of *Escherichia coli* B/r. J Bacteriol 119, 270–281 (1974).

9. Scott, M. & Hwa, T. Bacterial growth laws and their applications. Current Opinion in Biotechnology 22, 559–565 (2011).

10. Matamouros, S. et al. Growth-rate dependency of ribosome abundance and translation elongation rate in Corynebacterium glutamicum differs from that in Escherichia coli. Nat Commun 14, 5611 (2023).

11. Wilson, D. N. Ribosome-targeting antibiotics and mechanisms of bacterial resistance. Nat Rev Microbiol 12, 35–48 (2014).

12. Blanchard, S. C., Cooperman, B. S. & Wilson, D. N. Probing Translation with Small-Molecule Inhibitors. Chemistry & Biology 17, 633–645 (2010).

13. Neidhardt, F. C. & Magasanik, B. Studies on the role of ribonucleic acid in the growth of bacteria. Biochimica et Biophysica Acta 42, 99–116 (1960).

14. Mori, M. et al. From coarse to fine: the absolute Escherichia coli proteome under diverse growth conditions. Mol Syst Biol 17, MSB20209536 (2021).

15. Hui, S. et al. Quantitative proteomic analysis reveals a simple strategy of global resource allocation in bacteria. Mol Syst Biol 11, MSB145697 (2015).

16. Basan, M. et al. Overflow metabolism in Escherichia coli results from efficient proteome allocation. Nature 528, 99–104 (2015).

17. Deris, J. B. et al. The Innate Growth Bistability and Fitness Landscapes of Antibiotic-Resistant Bacteria. Science 342, 1237435 (2013).

18. Elf, J., Nilsson, K., Tenson, T. & Ehrenberg, M. Bistable Bacterial Growth Rate in Response to Antibiotics with Low Membrane Permeability. Phys. Rev. Lett. 97, 258104 (2006).

19. Frenkel, N., Saar Dover, R., Titon, E., Shai, Y. & Rom-Kedar, V. Bistable Bacterial Growth Dynamics in the Presence of Antimicrobial Agents. Antibiotics 10, 87 (2021).

20. Loewe, S. Die quantitativen Probleme der Pharmakologie. Ergebnisse der Physiologie 27, 47–187 (1928).

21. Bollenbach, T. Antimicrobial interactions: mechanisms and implications for drug discovery and resistance evolution. Current Opinion in Microbiology 27, 1–9 (2015).

22. Yeh, P., Tschumi, A. I. & Kishony, R. Functional classification of drugs by properties of their pairwise interactions. Nat Genet 38, 489–494 (2006).

23. Roemhild, R., Bollenbach, T. & Andersson, D. I. The physiology and genetics of bacterial responses to antibiotic combinations. Nat Rev Microbiol 20, 478–490 (2022).

24. Lázár, V., Snitser, O., Barkan, D. & Kishony, R. Antibiotic combinations reduce Staphylococcus aureus clearance. Nature 610, 540–546 (2022).

25. Fatsis-Kavalopoulos, N., Roemhild, R., Tang, P.-C., Kreuger, J. & Andersson, D. I. CombiANT: Antibiotic interaction testing made easy. PLoS Biol 18, e3000856 (2020).

26. Brochado, A. R. et al. Species-specific activity of antibacterial drug combinations. Nature 559, 259–263 (2018).

27. Zhang, Q. et al. A Decrease in Transcription Capacity Limits Growth Rate upon Translation Inhibition. mSystems 5, e00575–20 (2020).

28. Balakrishnan, R. et al. Principles of gene regulation quantitatively connect DNA to RNA and proteins in bacteria. Science 378, eabk2066 (2022).

29. Wu, C. et al. Cellular perception of growth rate and the mechanistic origin of bacterial growth law. Proc. Natl. Acad. Sci. U.S.A. 119, e2201585119 (2022).

30. Calabrese, L., Ciandrini, L. & Cosentino Lagomarsino, M. How total mRNA influences cell growth. Proc. Natl. Acad. Sci. U.S.A. 121, e2400679121 (2024).

31. Espinosa, R., Sørensen, M. A. & Svenningsen, S. L. Escherichia coli protein synthesis is limited by mRNA availability rather than ribosomal capacity during phosphate starvation. Front. Microbiol. 13, 989818 (2022).

32. Campbell, E. A. et al. Structural Mechanism for Rifampicin Inhibition of Bacterial RNA Polymerase. Cell 104, 901–912 (2001).

33. Si, F. et al. Invariance of Initiation Mass and Predictability of Cell Size in Escherichia coli. Current Biology 27, 1278–1287 (2017).

34. Roy, A., Goberman, D. & Pugatch, R. A unifying autocatalytic network-based framework for bacterial growth laws. Proc Natl Acad Sci U S A 118, e2107829118 (2021).

35. Geva-Zatorsky, N. et al. Protein Dynamics in Drug Combinations: a Linear Superposition of Individual-Drug Responses. Cell 140, 643–651 (2010).

36. Bollenbach, T. & Kishony, R. Resolution of Gene Regulatory Conflicts Caused by Combinations of Antibiotics. Molecular Cell 42, 413–425 (2011).

37. Lukačišin, M. & Bollenbach, T. Emergent Gene Expression Responses to Drug Combinations Predict Higher-Order Drug Interactions. Cell Systems 9, 423–433.e3 (2019).

38. Russo, C. J., Husain, K. & Murugan, A. Soft Modes as a Predictive Framework for Low-Dimensional Biological Systems Across Scales. Annual Review of Biophysics 54, 401– 426 (2025).

39. Li, G.-W., Burkhardt, D., Gross, C. & Weissman, J. S. Quantifying Absolute Protein Synthesis Rates Reveals Principles Underlying Allocation of Cellular Resources. Cell 157, 624–635 (2014).

40. Zhu, M., Mori, M., Hwa, T. & Dai, X. Distantly related bacteria share a rigid proteome allocation strategy with flexible enzyme kinetics. Proc. Natl. Acad. Sci. U.S.A. 122, e2427091122 (2025).

41. Dai, X. et al. Reduction of translating ribosomes enables Escherichia coli to maintain elongation rates during slow growth. Nat Microbiol 2, 16231 (2016).

42. Tritton, T. R. Ribosome-tetracycline interactions. Biochemistry 16, 4133–4138 (1977).

43. Davies, J. & Davis, B. D. Misreading of Ribonucleic Acid Code Words Induced by Aminoglycoside Antibiotics. Journal of Biological Chemistry 243, 3312–3316 (1968).

44. Schluenzen, F. et al. The antibiotic kasugamycin mimics mRNA nucleotides to destabilize tRNA binding and inhibit canonical translation initiation. Nat Struct Mol Biol 13, 871–878 (2006).

45. Eaton, D. S. et al. Essentialome-Wide Multigenerational Imaging Reveals Mechanistic Origins of Cell Growth Laws. Preprint at 10.1101/2025.06.10.658525 (2025).

46. Zhu, M., Mori, M., Hwa, T. & Dai, X. Disruption of transcription–translation coordination in Escherichia coli leads to premature transcriptional termination. Nat Microbiol 4, 2347–2356 (2019).

47. Karp, P. D. et al. The EcoCyc database (2025). EcoSal Plus 13, eesp-0019-2024 (2025).

48. Qi, Q., Angermayr, S. A. & Bollenbach, T. Uncovering Key Metabolic Determinants of the Drug Interactions Between Trimethoprim and Erythromycin in Escherichia coli. Front. Microbiol. 12, 760017 (2021).

49. Keseler, I. M. et al. EcoCyc: A comprehensive view of Escherichia coli biology. Nucleic Acids Research 37, D464–D470 (2009).

50. Hauryliuk, V., Atkinson, G. C., Murakami, K. S., Tenson, T. & Gerdes, K. Recent functional insights into the role of (p)ppGpp in bacterial physiology. Nat Rev Microbiol 13, 298–309 (2015).

51. Irving, S. E., Choudhury, N. R. & Corrigan, R. M. The stringent response and physiological roles of (pp)pGpp in bacteria. Nat Rev Microbiol 19, 256–271 (2021).

52. Potrykus, K. & Cashel, M. (p)ppGpp: Still Magical? Annu. Rev. Microbiol. 62, 35–51 (2008).

53. Ripamonti, A., Lacassin, M., Droghetti, R., Bokinsky, G. & Cosentino Lagomarsino, M. Transcriptional competition biases the effects of second messengers in Escherichia coli. Cell Systems 17, 101609 (2026).

54. Tuske, S. et al. Inhibition of Bacterial RNA Polymerase by Streptolydigin: Stabilization of a Straight-Bridge-Helix Active-Center Conformation. Cell 122, 541–552 (2005).

55. Temiakov, D. et al. Structural Basis of Transcription Inhibition by Antibiotic Streptolydigin. Molecular Cell 19, 655–666 (2005).

56. Hughes, J. & Mellows, G. Inhibition of Isoleucyl-Transfer Ribonucleic Acid Synthetase in Escherichia coli by Pseudomonic Acid. 176, (1978).

57. Nathans, D. PUROMYCIN INHIBITION OF PROTEIN SYNTHESIS: INCORPORATION OF PUROMYCIN INTO PEPTIDE CHAINS. Proc. Natl. Acad. Sci. U.S.A. 51, 585–592 (1964).

58. Thiermann, R. et al. Decoupling of global metabolic flux and proteome partitioning in bacteria. Science 392, eaeb6410 (2026).

59. Searle, B. C. et al. Generating high quality libraries for DIA MS with empirically corrected peptide predictions. Nat Commun 11, 1548 (2020).

60. Chambers, M. C. et al. A cross-platform toolkit for mass spectrometry and proteomics. Nat Biotechnol 30, 918–920 (2012).

61. Demichev, V., Messner, C. B., Vernardis, S. I., Lilley, K. S. & Ralser, M. DIA-NN: neural networks and interference correction enable deep proteome coverage in high throughput. Nat Methods 17, 41–44 (2020).

62. Perez-Riverol, Y. et al. The PRIDE database at 20 years: 2025 update. Nucleic Acids Res 53, D543–D553 (2025).

