## Supplementary Text for "A generalized growth law for translation- and transcription-targeting antibiotics captures drug interactions"

#### Extension of the growth law-based model of bacterial growth

This supplementary note describes an extension of the existing bacterial growth model based on growth laws [1–4], which we use to examine the effects of drug-specific ribosome responses on antibiotic susceptibility (Fig. S4) and drug interactions (Fig. 3). We extend the existing framework for ribosome-targeting antibiotics [2, 4] by allowing the ribosome–growth relation to vary among antibiotics. This difference is implemented via the phenomenological ribosome-response parameter $\theta$, which captures different ribosome reallocation responses caused by antibiotic perturbations (Fig. 1). In the single-drug case, changes in $\theta$ primarily rescale susceptibility through an effective ribosome range $\Delta r_\theta$ (Fig. S4A). In the two-drug case, one drug alters the physiological background in which the second drug acts, making susceptibility background-dependent and generating non-additive drug interactions (Fig. 3).

##### Mathematical model for individual translation inhibitors

In line with previous models of microbial resource reallocation [1–4], we assume that growth is determined by the unbound ribosome pool $r_\mathrm{u}$, which is related to the growth rate by the nutrient growth law

$$r_\mathrm{u}(\lambda) = r_\mathrm{min} + \frac{\lambda}{\kappa_\mathrm{t}}. \tag{1}$$

Here, $\lambda$ is the growth rate under the considered condition, $r_\mathrm{min}$ is the inactive non-growth-associated ribosome concentration and $\kappa_\mathrm{t}$ is the translational capacity. The drug-free growth rate in the same nutrient background is denoted by $\lambda_0$. Since no ribosomes are drug-bound in the drug-free state, the total ribosome concentration at this reference point is

$$r_0 \equiv r_\mathrm{tot}(\lambda_0) = r_\mathrm{min} + \frac{\lambda_0}{\kappa_\mathrm{t}}. \tag{2}$$

The total ribosome concentration is $r_\mathrm{tot} = r_\mathrm{u} + r_\mathrm{b}$, where $r_\mathrm{b}$ is the drug-bound ribosome concentration. The quantity $r_0$ denotes the total ribosome concentration in the drug-free reference state.

Following [2], the dynamics of antibiotics inhibiting translation are described by transport across the cell envelope and reversible binding to ribosomes. Drug transport into the cell is written as the flux

$$J(a_{\mathrm{ex}}, a) = p_{\mathrm{in}} a_{\mathrm{ex}} - p_{\mathrm{out}} a, \tag{3}$$

where $a_{\mathrm{ex}}$ and $a$ denote external and intracellular free drug concentrations, respectively. The parameters $p_{\mathrm{in}}$ and $p_{\mathrm{out}}$ describe effective influx and efflux permeabilities.
Drug binding to the growth-associated unbound ribosome pool is described by the net binding flux

$$f(r_{\mathrm{u}}, r_{\mathrm{b}}, a) = k_{\mathrm{on}} a \left(r_{\mathrm{u}} - r_{\mathrm{min}}\right) - k_{\mathrm{off}} r_{\mathrm{b}}, \tag{4}$$

where $k_{\mathrm{on}}$ and $k_{\mathrm{off}}$ are the binding and unbinding rates, respectively. The term $r_{\mathrm{u}} - r_{\mathrm{min}}$ represents the growth-associated unbound ribosome pool available for productive translation and drug binding. We assume that inactive ribosomes are not part of the ribosome pool accessible to the drug. In the model, $r_{\mathrm{min}}$ represents a ribosome fraction that does not contribute to productive translation. Changes in $r_{\mathrm{min}}$ between drugs are not explicitly represented.
We take dilution, drug transport and drug binding into account: The complete dynamical system reads:

$$\frac{\mathrm{d}a}{\mathrm{d}t} = -\lambda a - f(r_{\mathrm{u}}, r_{\mathrm{b}}, a) + J(a_{\mathrm{ex}}, a), \tag{5}$$

$$\frac{\mathrm{d}r_{\mathrm{u}}}{\mathrm{d}t} = -\lambda r_{\mathrm{u}} + s(\lambda) - f(r_{\mathrm{u}}, r_{\mathrm{b}}, a), \tag{6}$$

$$\frac{\mathrm{d}r_{\mathrm{b}}}{\mathrm{d}t} = -\lambda r_{\mathrm{b}} + f(r_{\mathrm{u}}, r_{\mathrm{b}}, a). \tag{7}$$

$s(\lambda)$ denotes the ribosome synthesis term. At steady state during balanced exponential growth, all time derivatives vanish and ribosome synthesis is given by $s(\lambda) = \lambda(r_{\mathrm{u}} + r_{\mathrm{b}})$.
With this sign convention, $f$ is defined as the net flux into the drug-bound ribosome pool. A positive value of $f$ means that free intracellular drug $a$ and unbound ribosomes $r_{\mathrm{u}}$ are consumed, while drug-bound ribosomes $r_{\mathrm{b}}$ are produced. The binding flux therefore enters with a negative sign in the equations for $a$ and $r_{\mathrm{u}}$ and with a positive sign in the equation for $r_{\mathrm{b}}$.
A full derivation of the steady-state solution can be found in [2]. We summarize the main steps relevant for the generalized model.
Starting from the steady-state condition for bound ribosomes, Eq. (7) gives

$$0 = -\lambda r_{\mathrm{b}} + f. \tag{8}$$

Combining this relation with the binding flux in Eq. (4) gives

$$\left(\lambda + k_{\mathrm{off}}\right) r_{\mathrm{b}} = k_{\mathrm{on}} a \left(r_{\mathrm{u}} - r_{\mathrm{min}}\right). \tag{9}$$

Using the nutrient growth law in Eq. (1), the steady-state amount of drug-bound ribosomes becomes

$$r_{\mathrm{b}} = \frac{k_{\mathrm{on}} a \lambda}{\kappa_{\mathrm{t}} \left(\lambda + k_{\mathrm{off}}\right)}. \tag{10}$$

Substituting Eq. (10) into the steady-state drug balance, Eq. (5), gives

$$0 = -\left(\lambda + p_{\mathrm{out}}\right) a - \lambda r_{\mathrm{b}} + p_{\mathrm{in}} a_{\mathrm{ex}}. \tag{11}$$

Using Eq. (10), this can be written as

$$p_{\mathrm{in}} a_{\mathrm{ex}} = a \left[ p_{\mathrm{out}} + \lambda + \frac{k_{\mathrm{on}} \lambda^2}{\kappa_{\mathrm{t}} \left(\lambda + k_{\mathrm{off}}\right)} \right]. \tag{12}$$

Thus, the intracellular drug concentration is

$$a = \frac{p_{\mathrm{in}} a_{\mathrm{ex}}}{p_{\mathrm{out}} + \lambda + \dfrac{k_{\mathrm{on}} \lambda^2}{\kappa_{\mathrm{t}} \left(\lambda + k_{\mathrm{off}}\right)}}. \tag{13}$$

To connect the drug-transport part of the model with the physiological ribosome balance, we first express the drug-bound ribosome concentration in terms of the external drug concentration and the growth rate. Substituting Eq. (13) into Eq. (10) gives

$$r_{\mathrm{b}}(a_{\mathrm{ex}}, \lambda) = \frac{k_{\mathrm{on}} p_{\mathrm{in}} a_{\mathrm{ex}} \lambda}{\kappa_{\mathrm{t}} \left(\lambda + k_{\mathrm{off}}\right) \left(p_{\mathrm{out}} + \lambda\right) + k_{\mathrm{on}} \lambda^2}. \tag{14}$$

This gives the kinetic expression for the amount of drug-bound ribosomes. In the next step, the same quantity is related to ribosome synthesis and growth through the steady-state ribosome balance.
The steady-state condition for unbound ribosomes, Eq. (6), is

$$0 = -\lambda r_{\mathrm{u}} + s(\lambda) - f. \tag{15}$$

Using Eq. (8), this becomes

$$s(\lambda) = \lambda \left(r_{\mathrm{u}} + r_{\mathrm{b}}\right) = \lambda r_{\mathrm{tot}}. \tag{16}$$

Equivalently,

$$s(\lambda) - \lambda r_{\mathrm{u}} = \lambda r_{\mathrm{b}}. \tag{17}$$

Using Eq. (1), this results in

$$s(\lambda) - \lambda r_{\mathrm{min}} - \frac{\lambda^2}{\kappa_{\mathrm{t}}} = \lambda r_{\mathrm{b}}. \tag{18}$$

Combining Eq. (18) with Eqs. (10) and (13) yields the compact steady-state identity

$$\left[ s(\lambda) - \lambda r_{\mathrm{min}} - \frac{\lambda^2}{\kappa_{\mathrm{t}}} \right] D(\lambda) = k_{\mathrm{on}} p_{\mathrm{in}} a_{\mathrm{ex}} \lambda^2, \tag{19}$$

with

$$D(\lambda) = \kappa_{\mathrm{t}}\,(\lambda + k_{\mathrm{off}})\,(p_{\mathrm{out}} + \lambda) + k_{\mathrm{on}}\lambda^2. \tag{20}$$

This equation forms the starting point for the generalized model. The kinetic contribution $D(\lambda)$, which contains transport and binding parameters, is separated from the physiological closure $s(\lambda)$, which describes how ribosome synthesis and proteome allocation change with growth rate. Drug kinetics and drug-induced proteome reorganization can be varied independently.

#### Generalized ribosome response for individual drugs

The established model assumes that total ribosome content increases linearly as the growth rate decreases due to translation inhibition [2]. More generally, different antibiotics induce distinct ribosome responses. While chloramphenicol treatment leads to strong compensatory ribosome upregulation, other perturbations, such as kasugamycin treatment or transcription limitation with rifampicin, produce substantially weaker or even opposite ribosome responses (Fig. 1). To account for these differences, we generalize the ribosome–growth relationship by introducing the phenomenological ribosome-response parameter $\theta$:

$$r_{\mathrm{tot}}(\lambda;\theta) = r_0 + \theta s_{\mathrm{G}}\,(\lambda - \lambda_0)\,. \tag{21}$$

Here, $r_0$ and $\lambda_0$ denote the drug-free reference state defined in Eq. (2). All generalized ribosome-response relations are required to pass through the same drug-free reference point $(\lambda_0, r_0)$. This ensures that changing $\theta$ changes the response to inhibition, but not the drug-free baseline.

The slope $s_{\mathrm{G}}$ is defined as the slope between the drug-free point $(\lambda_0, r_0)$ and the maximally inhibited ribosome state $r_{\max} = r_{\min} + \Delta r$, for chloramphenicol-like translation inhibition, as originally described [2],

$$\begin{aligned} s_{\mathrm{G}} &= \frac{r_{\max} - r_0}{0 - \lambda_0} \\ &= \frac{(r_{\min} + \Delta r) - (r_{\min} + \lambda_0/\kappa_{\mathrm{t}})}{-\lambda_0} \\ &= \frac{1}{\kappa_{\mathrm{t}}} - \frac{\Delta r}{\lambda_0}. \end{aligned} \tag{22}$$

Setting $\theta = 1$ recovers the previously reported response [2]. Varying $\theta$ alters the strength and direction of the ribosome reallocation response while preserving the same drug-free reference point $(\lambda_0, r_0)$. Here, $\theta = 1$ corresponds to the established ribosome increase of the second growth law, $0 < \theta < 1$ corresponds to a weaker increase, $\theta \approx 0$ corresponds to little or no drug-induced change in total ribosome content and $\theta < 0$ corresponds to a decrease in ribosome concentration in response to a drug.

Using Eq. (16), the synthesis term for the generalized closure is

$$s(\lambda) = \lambda r_{\mathrm{tot}}(\lambda;\theta) = \lambda\left[r_0 + \theta s_{\mathrm{G}}(\lambda - \lambda_0)\right]. \tag{23}$$

The physiological part of the steady-state identity is

$$B(\lambda) := s(\lambda) - \lambda r_{\min} - \frac{\lambda^2}{\kappa_{\mathrm{t}}}. \tag{24}$$

Substituting Eq. (23) and applying $r_0 - r_{\min} = \lambda_0/\kappa_{\mathrm{t}}$ yields

$$\begin{aligned} B(\lambda) &= \lambda r_0 + \theta s_{\mathrm{G}}\lambda(\lambda - \lambda_0) - \lambda r_{\min} - \frac{\lambda^2}{\kappa_{\mathrm{t}}} \\ &= \frac{\lambda\lambda_0}{\kappa_{\mathrm{t}}} - \frac{\lambda^2}{\kappa_{\mathrm{t}}} + \theta s_{\mathrm{G}}\lambda(\lambda - \lambda_0) \\ &= \left(\frac{1}{\kappa_{\mathrm{t}}} - \theta s_{\mathrm{G}}\right)\lambda(\lambda_0 - \lambda). \end{aligned} \tag{25}$$

This motivates defining the effective ribosome range

$$\Delta r_\theta = \lambda_0\left(\frac{1}{\kappa_{\mathrm{t}}} - \theta s_{\mathrm{G}}\right). \tag{26}$$

With this definition, Eq. (25) can be written as

$$B(\lambda) = \frac{\Delta r_\theta}{\lambda_0}\lambda(\lambda_0 - \lambda). \tag{27}$$

Substituting Eq. (27) into the general steady-state identity gives

$$\frac{\Delta r_\theta}{\lambda_0}\lambda(\lambda_0 - \lambda)D(\lambda) = k_{\mathrm{on}}p_{\mathrm{in}}a_{\mathrm{ex}}\lambda^2. \tag{28}$$

For $\lambda > 0$, this becomes

$$(\lambda_0 - \lambda)D(\lambda) = \frac{k_{\mathrm{on}}p_{\mathrm{in}}a_{\mathrm{ex}}\lambda_0}{\Delta r_\theta}\lambda. \tag{29}$$

Thus, the response parameter $\theta$ enters through the effective ribosome range $\Delta r_\theta$, while the kinetic denominator $D(\lambda)$ remains unchanged.
At half-maximal growth, $\lambda = \lambda_0/2$, Eq. (29) gives

$$\mathrm{IC}_{50,\theta} = \frac{\Delta r_\theta}{k_{\mathrm{on}}p_{\mathrm{in}}\lambda_0}D\left(\frac{\lambda_0}{2}\right). \tag{30}$$

Relative to the previously reported case,

$$\mathrm{IC}_{50,\theta} = \mathrm{IC}_{50,\mathrm{G}}\,\frac{\Delta r_\theta}{\Delta r}. \tag{31}$$

This shows that changing $\theta$ shifts susceptibility through proteome reorganization, while changes in dose-response shape require altered kinetic parameters, such as drug binding, unbinding, or transport. The generalized model remains physically meaningful only if

$$\Delta r_\theta > 0. \tag{32}$$

Strongly negative values of $\theta$ can violate this condition and lead to nonphysical solutions.

#### Generalized ribosome response for combinations of two drugs

Next, we extend the model to a drug combination setting, in which one drug modifies the physiological background and the other drug acts within this altered state. We use a reduced model for this conditional response. In the main orientation used below, drug X defines the background and drug Y is evaluated in this background. Rifampicin is an example of drug X. It is represented through its single-drug growth response and is not assumed to follow the same ribosome-binding kinetics as the translation inhibitors in the one-drug model. We represent the physiological state induced by the background drug by its growth rate and ribosome state. Other drug-induced changes in cellular physiology are not modeled separately.

The single-drug growth rate in the background generated by drug X is written as

$$\lambda_0(c_{\mathrm{X}}) = \lambda_{00} g_{\mathrm{X}}(c_{\mathrm{X}}), \qquad g_{\mathrm{X}}(0) = 1. \tag{33}$$

Here, $c_{\mathrm{X}}$ is the concentration of the background drug, $\lambda_{00}$ is the growth rate in the absence of both drugs and $g_{\mathrm{X}}(c_{\mathrm{X}})$ is the normalized single-drug growth response of drug X.

We approximate the corresponding background ribosome level using the nutrient growth-law relation,

$$r_0(c_{\mathrm{X}}) = r_{\min} + \frac{\lambda_0(c_{\mathrm{X}})}{\kappa_{\mathrm{t}}}. \tag{34}$$

This is a modeling assumption for the X-only background state. It does not imply that the complete two-drug response follows the nutrient growth-law line. The corresponding background-dependent slope is

$$s_{\mathrm{G}}(c_{\mathrm{X}}) = \frac{1}{\kappa_{\mathrm{t}}} - \frac{\Delta r}{\lambda_0(c_{\mathrm{X}})}. \tag{35}$$

The effect of drug Y is evaluated within this X-dependent physiological state according to

$$r_{\mathrm{tot}}\left(\lambda; c_{\mathrm{X}}, \theta_{\mathrm{Y}}\right) = r_0(c_{\mathrm{X}}) + \theta_{\mathrm{Y}} s_{\mathrm{G}}(c_{\mathrm{X}}) \left[\lambda - \lambda_0(c_{\mathrm{X}})\right]. \tag{36}$$

Here, $\theta_{\mathrm{Y}}$ is the ribosome-response parameter of the drug whose response is evaluated in the background. In this reduced model, $\theta_{\mathrm{Y}}$ is treated as a drug-specific constant with respect to the background concentration $c_{\mathrm{X}}$. This assumption could fail if the response class of drug Y changes strongly across different X-dependent backgrounds.

The effective susceptibility of drug Y therefore depends on the background through the scale $\lambda_0(c_{\mathrm{X}})/\Delta r_{\theta,\mathrm{Y}|\mathrm{X}}(c_{\mathrm{X}})$. The effective ribosome range associated with the response to drug Y is

$$\Delta r_{\theta,\mathrm{Y}|\mathrm{X}}(c_{\mathrm{X}}) = \lambda_0(c_{\mathrm{X}}) \left[\frac{1}{\kappa_{\mathrm{t}}} - \theta_{\mathrm{Y}} s_{\mathrm{G}}(c_{\mathrm{X}})\right]. \tag{37}$$

Using Eq. (35), this can be written as

$$\begin{aligned}\Delta r_{\theta,\mathrm{Y|X}}(c_{\mathrm{X}}) &= \lambda_0(c_{\mathrm{X}})\left[\frac{1}{\kappa_{\mathrm{t}}} - \theta_{\mathrm{Y}}\left(\frac{1}{\kappa_{\mathrm{t}}} - \frac{\Delta r}{\lambda_0(c_{\mathrm{X}})}\right)\right] \\ &= (1-\theta_{\mathrm{Y}})\frac{\lambda_0(c_{\mathrm{X}})}{\kappa_{\mathrm{t}}} + \theta_{\mathrm{Y}}\Delta r.\end{aligned} \tag{38}$$

The same construction can be written after exchanging the two drugs. Drug Y then defines the background and drug X acts in this background. This gives

$$\Delta r_{\theta,\mathrm{X|Y}}(c_{\mathrm{Y}}) = (1-\theta_{\mathrm{X}})\frac{\lambda_0(c_{\mathrm{Y}})}{\kappa_{\mathrm{t}}} + \theta_{\mathrm{X}}\Delta r. \tag{39}$$

The algebra has the same form after exchanging $X$ and $Y$, but the response parameter belongs to the drug evaluated in the background. The $Y|X$ direction depends on $\theta_{\mathrm{Y}}$, while the $X|Y$ direction depends on $\theta_{\mathrm{X}}$. The two conditional descriptions need not give the same response.

We continue with the $X \to Y$ orientation, where drug X defines the background and drug Y is evaluated in this background. In the one-drug case, $\Delta r_\theta$ is constant for a given response class. In the two-drug case, the physiological background created by drug X changes the effective susceptibility scale of drug Y across the $c_{\mathrm{X}}$-axis. Each fixed concentration of drug X defines a different effective one-dimensional dose-response curve for drug Y. These conditional dose-response curves form a two-dimensional response surface for the combination of the two drugs (Fig. 3).

The simple background closure in Eq. (34) assumes that the background ribosome level lies on the nutrient growth-law line. If the background drug changes ribosome allocation independently of this line, the background state could be written more generally as

$$R_{\mathrm{X}}(c_{\mathrm{X}}) = r_{\min} + \frac{\lambda_0(c_{\mathrm{X}})}{\kappa_{\mathrm{t}}} + \varepsilon_{\mathrm{X}}(c_{\mathrm{X}}), \tag{40}$$

where $\varepsilon_{\mathrm{X}}(c_{\mathrm{X}})$ describes the deviation of the X-only background from the nutrient growth-law line. The reduced model used here corresponds to the special case $\varepsilon_{\mathrm{X}}(c_{\mathrm{X}}) = 0$. This becomes important when the background perturbation is rifampicin-like rather than nutrient-like.

For each fixed background concentration $c_{\mathrm{X}}$, the response to drug Y follows the same form as Eq. (29):

$$[\lambda_0(c_{\mathrm{X}}) - \lambda]\, D_{\mathrm{Y}}(\lambda) = \frac{k_{\mathrm{on,Y}}\, p_{\mathrm{in,Y}}\, c_{\mathrm{Y}}\, \lambda_0(c_{\mathrm{X}})}{\Delta r_{\theta,\mathrm{Y|X}}(c_{\mathrm{X}})}\lambda, \tag{41}$$

with

$$D_{\mathrm{Y}}(\lambda) = \kappa_{\mathrm{t}}\,(\lambda + k_{\mathrm{off,Y}})\,(p_{\mathrm{out,Y}} + \lambda) + k_{\mathrm{on,Y}}\lambda^2. \tag{42}$$

Here, $c_{\mathrm{Y}}$ is the external concentration of drug Y and $k_{\mathrm{on,Y}}$, $k_{\mathrm{off,Y}}$, $p_{\mathrm{in,Y}}$ and $p_{\mathrm{out,Y}}$ are the kinetic parameters of drug Y.

Changes in the effective susceptibility scale

$$\lambda_0(c_X)/\Delta r_{\theta,Y|X}(c_X)$$

with the background concentration $c_X$ can contribute to curvature of the drug interaction surface. Part of this dependence comes from changes in the effective ribosome range. Differentiating Eq. (37) with respect to $c_X$ gives

$$\frac{d\Delta r_{\theta,Y|X}(c_X)}{dc_X} = \lambda_0'(c_X)\left[\frac{1}{\kappa_t} - \theta_Y s_G(c_X)\right] - \lambda_0(c_X)\theta_Y s_G'(c_X). \tag{43}$$

The first term reflects the change in background growth rate caused by drug X. The second term reflects the change in the growth-law slope and is weighted by the proteome-response class $\theta_Y$.

Using the expanded expression in Eq. (38), the $Y|X$ susceptibility scale can be written as

$$S_{Y|X}(c_X) = \frac{\lambda_0(c_X)}{\Delta r_{\theta,Y|X}(c_X)} = \frac{\lambda_0(c_X)}{(1-\theta_Y)\lambda_0(c_X)/\kappa_t + \theta_Y \Delta r}. \tag{44}$$

The two limiting cases illustrate how this background dependence changes with $\theta_Y$. For $\theta_Y = 1$, corresponding to a strong compensatory ribosome response, the effective ribosome range becomes

$$\Delta r_{\theta,Y|X}(c_X) = \Delta r, \tag{45}$$

Therefore

$$S_{Y|X}(c_X) = \frac{\lambda_0(c_X)}{\Delta r}. \tag{46}$$

Thus, for $\theta_Y = 1$, the susceptibility scale remains directly proportional to the background growth rate $\lambda_0(c_X)$.

For $\theta_Y = 0$, corresponding to little or no additional drug-induced ribosome reallocation by drug Y, the effective ribosome range becomes

$$\Delta r_{\theta,Y|X}(c_X) = \frac{\lambda_0(c_X)}{\kappa_t}, \tag{47}$$

and therefore

$$S_{Y|X}(c_X) = \frac{\lambda_0(c_X)}{\lambda_0(c_X)/\kappa_t} = \kappa_t. \tag{48}$$

Thus, for $\theta_Y = 0$, the explicit $\lambda_0(c_X)$ factor cancels from the $Y|X$ susceptibility scale. The response of drug Y therefore no longer depends on the background growth rate through this term.

The same cancellation argument applies after exchanging the drugs. In the $X|Y$ direction, the corresponding susceptibility scale is

$$S_{X|Y}(c_Y) = \frac{\lambda_0(c_Y)}{\Delta r_{\theta,X|Y}(c_Y)}. \tag{49}$$

The analogous cancellation occurs only if $\theta_{\mathrm{X}} = 0$:

$$\Delta r_{\theta,\mathrm{X}|\mathrm{Y}}(c_{\mathrm{Y}}) = \frac{\lambda_0(c_{\mathrm{Y}})}{\kappa_{\mathrm{t}}} \quad \Rightarrow \quad S_{\mathrm{X}|\mathrm{Y}}(c_{\mathrm{Y}}) = \kappa_{\mathrm{t}}. \tag{50}$$

A value of $\theta_{\mathrm{Y}} = 0$ does not by itself imply that the conditional $Y|X$ response is additive. It only removes one source of background-growth dependence in this direction. After exchanging the drugs, the corresponding result depends on $\theta_{\mathrm{X}}$.

##### Drug-drug growth surface

A two-drug growth surface can be written as

$$\lambda = F(c_{\mathrm{X}}, c_{\mathrm{Y}}). \tag{51}$$

An isobole is a contour of equal growth,

$$F(c_{\mathrm{X}}, c_{\mathrm{Y}}) = \lambda^*. \tag{52}$$

Implicit differentiation along this contour gives

$$\frac{\mathrm{d}c_{\mathrm{Y}}}{\mathrm{d}c_{\mathrm{X}}} = -\frac{\partial F/\partial c_{\mathrm{X}}}{\partial F/\partial c_{\mathrm{Y}}}. \tag{53}$$

An isobole is curved when this ratio changes along the contour. In this reduced model, Eq. (53) describes the geometry of isobole curvature but does not provide a closed-form condition for additivity. The growth surface itself is obtained from the conditional dose-response equation, Eq. (41).

One source of curvature is the background-dependent susceptibility scale $S_{\mathrm{Y}|\mathrm{X}}(c_{\mathrm{X}})$. For $\theta_{\mathrm{Y}} = 1$, this scale remains proportional to the background growth rate $\lambda_0(c_{\mathrm{X}})$. For $\theta_{\mathrm{Y}} = 0$, the explicit $\lambda_0(c_{\mathrm{X}})$ factor cancels from this scale. This reduces one source of background dependence in the $Y|X$ direction.

This result applies to the $Y|X$ direction. In the exchanged $X|Y$ direction, the corresponding cancellation requires $\theta_{\mathrm{X}} = 0$. Thus, the two conditional descriptions need not give the same response.

For rifampicin–chloramphenicol, the model should therefore be interpreted phenomenologically. Rifampicin is used as drug X, which creates a transcription-limited background through $\lambda_0(c_{\mathrm{X}})$, while chloramphenicol acts as drug Y and is represented by a strong ribosome-upregulation response, $\theta_{\mathrm{Y}} \approx 1$. For rifampicin–kasugamycin, kasugamycin has a weaker ribosome reallocation response, so $\theta_{\mathrm{Y}} \approx 0$ is a plausible phenomenological description. In this reduced model, the kasugamycin response can therefore depend less strongly on the background than the chloramphenicol response.

Because the two drugs have different roles in this model, the response may depend on treatment order. The state generated by drug X before adding drug Y may differ from the state generated when the order is reversed. This could be tested experimentally by treating cells first with one drug and then with the other. The present steady-state model does not describe these time-dependent effects.

### References


1. Scott, M. & Hwa, T. Bacterial growth laws and their applications. en. *Current Opinion in Biotechnology* **22,** 559–565 (Aug. 2011).
2. Greulich, P., Scott, M., Evans, M. R. & Allen, R. J. Growth-dependent bacterial susceptibility to ribosome-targeting antibiotics. en. *Molecular Systems Biology* **11,** MSB145949 (Mar. 2015).
3. Kavčič, B., Tkačik, G. & Bollenbach, T. Mechanisms of drug interactions between translation-inhibiting antibiotics. en. *Nature Communications* **11,** 4013 (Aug. 2020).
4. Kavčič, B., Tkačik, G. & Bollenbach, T. Minimal biophysical model of combined antibiotic action. en. *PLOS Computational Biology* **17** (ed Grilli, J.) e1008529 (Jan. 2021).
